# Dynamic Allelic Expression in Mouse Mammary Gland Across the Adult Developmental Cycle

**DOI:** 10.1101/2024.09.02.610775

**Authors:** Geula Hanin, Kevin R Costello, Hugo Tavares, Boshra AlSulaiti, Shrina Patel, Carol A Edwards, Anne C Ferguson-Smith

**Author notes:** Corresponding author: Anne C Ferguson-Smith.

## Abstract

The mammary gland, which primarily develops postnatally, undergoes significant changes during pregnancy and lactation to facilitate milk production. Through the generation and analysis of 480 transcriptomes, we provide the most detailed allelic expression map of the mammary gland, cataloguing cell-type-specific expression from ex-vivo purified cell populations over 10 developmental stages, enabling comparative analysis. The work identifies genes involved in the mammary gland cycle, parental-origin-specific and genetic background-specific expression at cellular and temporal resolution, genes associated with human lactation disorders and breast cancer. Genomic imprinting, a mechanism regulating gene expression based on parental origin, is crucial for controlling gene dosage and stem cell potential throughout development. The analysis identified 25 imprinted genes monoallelically expressed in the mammary gland, with several showing allele-specific expression in distinct cell types. No novel imprinted genes were identified and the absence of biallelically expressed imprinted genes suggests that, unlike in brain, selective absence of imprinting does not regulate gene dosage in the mammary gland. This research highlights transcriptional dynamics within mammary gland cells and identifies novel candidate genes potentially significant in the tissue during gestation and lactation. Overall, this comprehensive atlas represents a valuable resource for future studies on expression and transcriptional dynamics in mammary cells.

## Introduction

The mammary gland is a unique organ that mostly develops postnatally in preparation to nourish the neonate. During pregnancy, in response to circulating hormones, the mammary gland undergoes dramatic growth, proliferation, and differentiation. This process is characterised by extensive ductal side branching and the formation of lobuloalveolar structures. By the end of pregnancy, the mammary gland is densely populated with alveoli, which become functional for milk production close to parturition^1,2^. Lactation marks a functional transition when the gland is fully dedicated to supplying the energetic demands of the offspring. Upon cessation of lactation, the mammary gland undergoes a tightly regulated involution process, during which the alveolar structures are removed, and the tissue remodels to a simpler ductal architecture^3^. While this structure resembles the pre-pregnancy state, the epigenetic landscape is modified^4^, priming the gland for subsequent pregnancies.

Various mammary gland cell types play crucial roles in this dynamic process. Mammary adipocytes form the mammary fat pad, providing structural support. Epithelial cells, comprised of basal and luminal cells, undergo dramatic changes during gestation, transforming into ductal cells, secretory alveolar cells and contractile myoepithelial cells. Other mammary cells are essential for proper function, including endothelial cells maintaining adequate blood supply, stromal cells such as fibroblasts and a variety of immune cells which are functional throughout the developmental cycle. The normal transcriptional dynamics and signalling processes involving these cell types at these stages are not well understood.

Genomic imprinting is a mammalian epigenetically regulated process causing genes to be expressed monoallelically according to their parental origin^5^. Genetic studies have demonstrated that genomic imprinting generally occurs at dosage-sensitive genes regulating critical developmental processes, with extensive focus on the embryo and placenta^6–8^. In humans, altered imprinted gene expression is associated with growth, metabolic and neurological disorders, and cancer^5^.

Genomic imprinting in mammals evolved alongside viviparity and is observed in both eutherian and metatherian mammals, but not, to date, in oviparous monotreme mammals^9^. The prevailing theory of imprinting evolution suggests it results from a conflict between maternally and paternally-derived genes in the conceptus, particularly around resource control, with a focus on the feto-placental interface and prenatal growth^10–13^. For example, deletion of the paternally expressed *Igf2* gene in the placenta reduces nutrient supply to the fetus, causing prenatal growth retardation^14,15^. In contrast, deletion of the maternally expressed *Grb10* gene leads to placental overgrowth and increased efficiency^16^. The imprinting of these genes suggests a parental tug-of-war over prenatal resource control as evidenced by the expression and imprinting of most prenatally transcribed imprinted genes in the placenta of mice.

In marsupials, with a shorter pregnancy and major nutritional provision occurring postnatally through lactation, the mammary gland plays a critical role in delivering milk to the developing neonate. Although fewer genes are imprinted in marsupials than in eutherians^17,18^, the mammary gland has been proposed as a preferred site for imprinted gene expression^19^. Specific imprinting of insulin (*Ins*) and *Igf2* in the marsupial mammary gland has been identified, suggesting a parallel to imprinting in the eutherian placenta^20^.

Despite several studies indicating roles for specific imprinted genes in the mammary gland (e.g., *Grb10*, *Meg3*, *Igf2*, *Peg1/Mest*)^16,21–23^, and similarities in imprinted gene expression patterns between the placenta and mammary gland^24^, the exact role of imprinting during lactation remains unclear. Previous reports lacked the resolution for a thorough exploration of imprinting in the mammary gland. Our previous systemic exploration of published work revealed numerous genes expressed in discrete mammary cell populations, some monoallelically^13^. Subsequent studies supported these findings but were limited by sample size and timing^25^.

To further understand imprinting in the mammary gland, it is important to determine its role in eutherian mammals. Although studies have been limited, *Igf2* has been shown to mediate prolactin-induced morphogenesis in the breast^23^, and *Grb10* has been shown to control milk supply and demand^16^. Contradictory findings exist for *Peg3*, with some studies showing its deletion impairs milk let-down, while others report no effect on nursing^21,26^. More recently, *Peg1/Mest* has also been implicated in mammary gland maturation in vitro^22^.

Interestingly, evidence suggests that imprinted genes can be selectively regulated in specific cell types or developmental time points as a mechanism to modulate gene dosage essential for normal development, particularly in stem cell populations. For example, the vertebrate-specific atypical Notch ligand, *Dlk1* (Delta-like homologue 1), which is usually expressed from the paternally inherited chromosome, shows selective absence of imprinting in the postnatal neurogenic niche, resulting in the activation of the canonically repressed maternal allele^27^. This biallelic expression of *Dlk1* is crucial for normal adult neurogenesis, as evidenced by premature exit from quiescence and depletion of neural stem cells in both maternal and paternal heterozygous *Dlk1* mutants. Similarly, *Igf2* exhibits selective biallelic expression in the choroid plexus to regulate neural stem cell proliferation^28,29^, underscoring the importance of precise dosage control by genomic imprinting in specific contexts. This raises the question of whether this is a common feature of stem cells *in vivo*, or is specific to the developing brain^30^.

Although these reports support the mammary gland as a site of imprinting, a comprehensive cellular and temporal dataset for in-depth exploration is lacking. Notably, since the father’s genome is not present in the mammary gland that is supporting the nutritional supply to his offspring, it challenges the theory of parental conflict over resource allocation. If imprinting is found to be important in the eutherian mammary gland, it might support alternative evolutionary theories, such as the maternal-offspring co-adaptation theory^31^. Understanding the role of imprinting in mammary gland development and function could provide significant insights into the developmental biology and the evolution of genomic imprinting.

Here we provide the first mammary gland cell type, stage and parent-of-origin-specific expression atlas. Our data from 480 transcriptomes identifies imprinted genes, expressed in the mammary gland in a monoallelic fashion. At the same time, our allelic atlas reveals that monoallelic expression of imprinted genes is not restricted to a small subset of genes, but rather a wider phenomenon. Therefore, it informs our understanding of the evolutionary origin of imprinting supporting a maternal-offspring coadaptation model for the evolution of imprinting.

## Materials and methods

### Animals

Mouse work and the experiments for this study were approved by the University of Cambridge Animal Welfare, and Ethical Review Body and conducted under the UK Home ORice Animals (Scientific Procedures) Act 1986. (Home ORice project licence # PC213320E and PP8193772).

Reciprocal hybrid mice were generated by crossing C57BL/6J females × CAST/EiJ males to produce BC hybrids or CAST/EiJ females × C57BL/6J males to produce CB hybrids. Mice were housed under a 12h light / 12h dark photocycle 22°C air temperature and 21% oxygen saturation with access to water and standard laboratory chow diet ad libitum (SAFE Diet 105, Safe lab, UK, containing 73.6% starch,19.3% protein, 5.1% fat, 3.3% simple sugars of energy contribution).

BC and CB F1 hybrid females were mated at 12-16 weeks of age, and mammary glands were collected at gestation day 5.5, 9.5 or 14.5, lactation days 5, 10 or 15 or involution days 1, 6 or 14, together creating 10 different time points throughout the developmental cycle of the adult mammary gland. Nulliparous mice were oestrus stage matched using Toludine Blue staining of vaginal smears, and collected at Oestrus. Female hybrids designated for gestation days were mated, with the day a vaginal plug was found designated as day 0.5, pregnancy was confirmed during dissections. Lactation days were counted from the day following parturition, with the first day designated lactation day 1. Involution days were counted from the removal of pups at postnatal day 21, with the day after pup removal considered involution day 1. Each time point and reciprocal cross comprised n=4 mice.

### Carmine staining

Whole mounts of mammary tissues from nulliparous mice, gestation and lactation stages we fixed in 4% Paraformaldehyde at 4°C, rinsed in Phosphate-buffered saline and stained overnight with carmine alum solution (Sigma-Aldrich C-1022, Aluminum Potassium sulphate (Sigma Aldrich A-7167). Whole mounts were dehydrated and mounted for imaging using Leica MZ16F.

### Flow cytometry

Single-cell suspensions of mammary cells were prepared from hybrid BC or CB females. Mammary glands were dissected and digested as previously described^32^. Mammary adipocytes were processed simultaneously, as previously described^33^.

Briefly, mammary fragments were digested for 1 hour at 37**°**C using Krebs–Ringers – HEPES, 2.5 mM glucose (Sigma-Aldrich, D9434), 2% fetal bovine serum (Gibco, 16-000-044), 200 μM adenosine (Sigma-Aldrich, A9251), 1 mg/mL collagenase (Sigma-Aldrich, C2139), pH=7.4. Further dissociation was performed using Trypsin-EDTA 0.25% (Gibco 25200072). Single cells were then filtered, and separated into the mammary adipocyte layer which was visible as the upper phase following centrifugation, and mammary cells which formed a pellet.

Cells were further incubated in 1mg/mL DNAse (Sigma Aldrich D4513), 5 mg/mL Dispase (Sigma Aldrich D4693) and 0.25% Trypsin EDTA (Thermo Fisher Scientific 25200056) to ensure a single cell suspension.

Cells were then blocked with 10% normal rat serum (Sigma-Aldrich R9759-10ML) for 15 minutes at 4°C. Pelleted mammary cells stained with the following antibodies: Anti-Mouse CD45 APC-eFluor780 (Invitrogen, 47-0451-82), Anti-Mouse CD31 PE-Cy7 (Invitrogen, 25-0311-82), Anti-Mouse TER-119 Biotin (Invitrogen, 13-5921-82), Anti-Mouse BP-1 Biotin (Invitrogen, 13-5891-81), Anti-Mouse EpCam Alexa Fluor647 (Biolegend, 118211), Anti-Mouse CD49b PE (Biolegend, 103506), Anti-Mouse CD49f Alexa Fluor488 (Biolegend, 313608). Viability was assessed using Zombie Aqua dye (Biolegend 423101).

Mammary adipocytes were stained with LipidTox Deep Red (Thermo Fischer, H34477) and the following antibodies were used to exclude other cell populations:CD45 biotin (Thermo Fischer, 13-0451-85), CD31 biotin (Thermo Fischer, 13-0311-85), CD49f Alexa Fluor 488 (AF488) (BioLegend 313608), Ter119 biotin (Thermo Fischer, 13-5921-82), BP-1 biotin (Thermo Fischer 13-5891-81), and Alexa Fluor 488 (AF488) Steptavidin (BioLegend 405235).

Cell populations were sorted using BD ARIA III cell sorter (BD Biosciences). Data were analysed using FlowJo 10.8.1.

### RNA extraction

RNA was extracted from sorted cells or whole mammary gland tissue using miRNeasy micro kit (QIAGEN, 217084), according to standard protocols. RNA concentration was determined using Qubit Fluorometer (ThermoFisher), and RNA integrity was quantified using the 2100 Bioanalyzer instrument (Agilent).

### Library generation and RNA sequencing

Libraries for RNA sequencing (RNA-seq) from sorted mammary cells populations including basal, stromal, luminal differentiated, luminal progenitors, endothelial cells and mammary adipocytes, were generated using SMARTer stranded total RNA-seq kit V2 – Pico input mammalian (TaKaRa bio, 634414) according to the manufacturer’s instructions using 13 cycles of amplification. The quality and RNA integrity number were assessed using Bioanalyzer 2100 (Agilent), Qubit fluorometer (Thermo Fisher) and sequenced using the NovaSeq 6000 system (Illumina), following a validation of index balance using miSeq (Illumina).

RNA-seq processing pipeline was built using Snakemake v6.10.0^34^. Reads were trimmed using Cutadapt v3.4^35^ with the following settings: Illumina TruSeq adapters, -U4 -- quality-cutoR 20, minimum length 50 base pairs. Read quality was assessed before and after trimming with FastQC v0.11.9 (Andrews 2010, http://www.bioinformatics.babraham.ac.uk/projects/fastqc). Further quality control metrics were obtained by aligning the reads to the standard GRCm38/mm10 mouse reference genome (Ensembl release 102) using STAR v2.7.9a^36^, followed by RSeQC v4.0.0^37^ for quality alignment control. Quality metrics were compiled using MultiQC v1.10.1^38^. Transcript quantification was performed using Salmon v1.5.0 in mapping-based mode^39^. Read count matrices were prepared using the R/Bioconductor package tximport v1.20.0^40^. DESeq2 v1.32.0^41^ was used to detect differentially expressed protein-coding genes with at least 20 reads, false discovery rate (FDR) threshold of 0.05 and an absolute log2 fold-change threshold of 1.0. Differential expression gene analysis was performed separately for each cell type and stage. The DESeq2 package was also used to obtain normalised read counts using the variance stabilising transformation method.

### Bioinformatic analysis

Allele-specific transcript quantification was processed by building a “diploid” reference genome based on the CAST/EiJ strain variants (SNPs and indels) from the mouse genome project (https://www.sanger.ac.uk/data/mouse-genomes-project/, using *bcftools consensus* v1.12. We then created a CAST/EiJ-specific transcript annotation GTF file using *STAR* v2.7.9a with ‘--runMode liftOver’, and extracted fasta files for each transcript using *gffread* v0.12.1. This resulted in two strain-specific transcript FASTA sequences, which were used to create “diploid” genome/transcriptome references. These were used to quantify allele-specific transcript abundance using *salmon* v1.5.0. The output of this pipeline allowed us to obtain both gene-level and transcript-level read counts either summarised across both alleles (for standard expression analysis) or individually for each allele (for allele-specific analysis). Read count matrices were prepared using the R/Bioconductor package *tximport* v1.20.0. Standard expression analysis was performed using the R/Bioconductor package *DESeq2* v1.32.0, in particular to obtain normalised read counts using the “variance stabilising transformation” method, which accounts for library composition differences as well as effective transcript length (as estimated by Salmon). Allele-specific expression analysis was performed using the R/Bioconductor package *ISoLDE* v1.20.0. The full pipeline was implemented in an automated workflow using *snakemake* v6.10.0. Bioinformatic pipeline used above for the mammary hybrid dataset is available at GitHub: https://github.com/AFS-lab/allele_specific_rnaseq_workflow Instructions are included in the repository.

### Profiling of GWAS targets

All single nucleotide polymorphisms (SNPs) associated with the terms “Breast milk measurement” and “Breastfeeding duration” were retrieved from the NHGRI-EBI GWAS catalogue to identify transcripts involved in the regulation of human lactation. SNPs related to human milk oligosaccharides were excluded from analysis due to their absence in the mouse genome. Transcripts impacted by these SNPs were further filtered to include only those with an RPKM > 1 in at least one profiled cell type or timepoint. Differentially expressed transcripts were selected using a Padj value less than 0.05 between the timepoint of interest relative to nulliparous. Standard expression analysis was performed using the R/Bioconductor package *DESeq2* v1.32.0, in particular to obtain normalised read counts using the “variance stabilising transformation” method, which accounts for library composition differences as well as effective transcript length (as estimated by Salmon). All SNPs identified in the breast disease category were selected to identify transcripts related to postpartum breast cancer. Transcripts found to be impacted by these SNPs were selected and filtered to only select for transcripts with an RPKM > 1 in at least one of the profiled cell types or timepoints. To determine all differentially expressed transcripts between the datasets, all transcripts with Padj value less than 0.05 between peak lactation, denoted as lactation d10 in this dataset, and all stages of involution were identified. Standard expression analysis was performed using the R/Bioconductor package *DESeq2* v1.32.0, in particular to obtain normalised read counts using the “variance stabilising transformation” method, which accounts for library composition differences as well as effective transcript length (as estimated by Salmon). To further select the most differentially expressed genes, non-protein-coding genes were excluded based on RefSeq annotations from the UCSC Genome Browser (https://pubmed.ncbi.nlm.nih.gov/39460617/). Transcripts showing the most significant differential expression (Padj < 10^−17^) across all stages of involution were selected. Average fold changes in expression for these transcripts were calculated and visualised, as shown in Figure 6C.

### qPCR

mRNA levels were determined by quantitative reverse transcription PCR (RT-qPCR) using the RevertAid H Minus First Strand cDNA Synthesis kit (Thermo Scientific) followed by quantification in technical triplicates with Brilliant III Ultra-Fast SYBR® Green QPCR Master Mix (Agilent) on the LightCycler 480 Instrument (Roche). Relative expression was normalised to β-Tubulin expression unless otherwise noted and calculated using the ΔCt method.

Primers were designed using Primer3 software. Primer sequences are listed in Supplementary Table 1

## Results

### Timelapse transcriptomes reveal cell-type-specific differentiation clustering, with lactation inducing the most significant changes beyond epithelial cells

To create a high-resolution atlas of adult cell-type-specific mammary gland allelic transcriptome, we generated a cohort of reciprocal hybrid mice with distinct parental origins. Specifically, we crossed CAST/EiJ female mice with C57Bl6/J males to produce F1 CB hybrid mice and conducted the reciprocal cross to generate BC hybrid mice (Figure 1A). To encompass the entire adult developmental cycle of the mammary gland and create a thorough transcriptome we selected ten key time points representing early, mid and late stages of mammary gland development: nulliparous stage, gestation days 5.5, 9.5 and 14.5, lactation days 5, 10 and 15 and involution days 1, 6 and 14 (Figure 1B).

**Figure 1.**
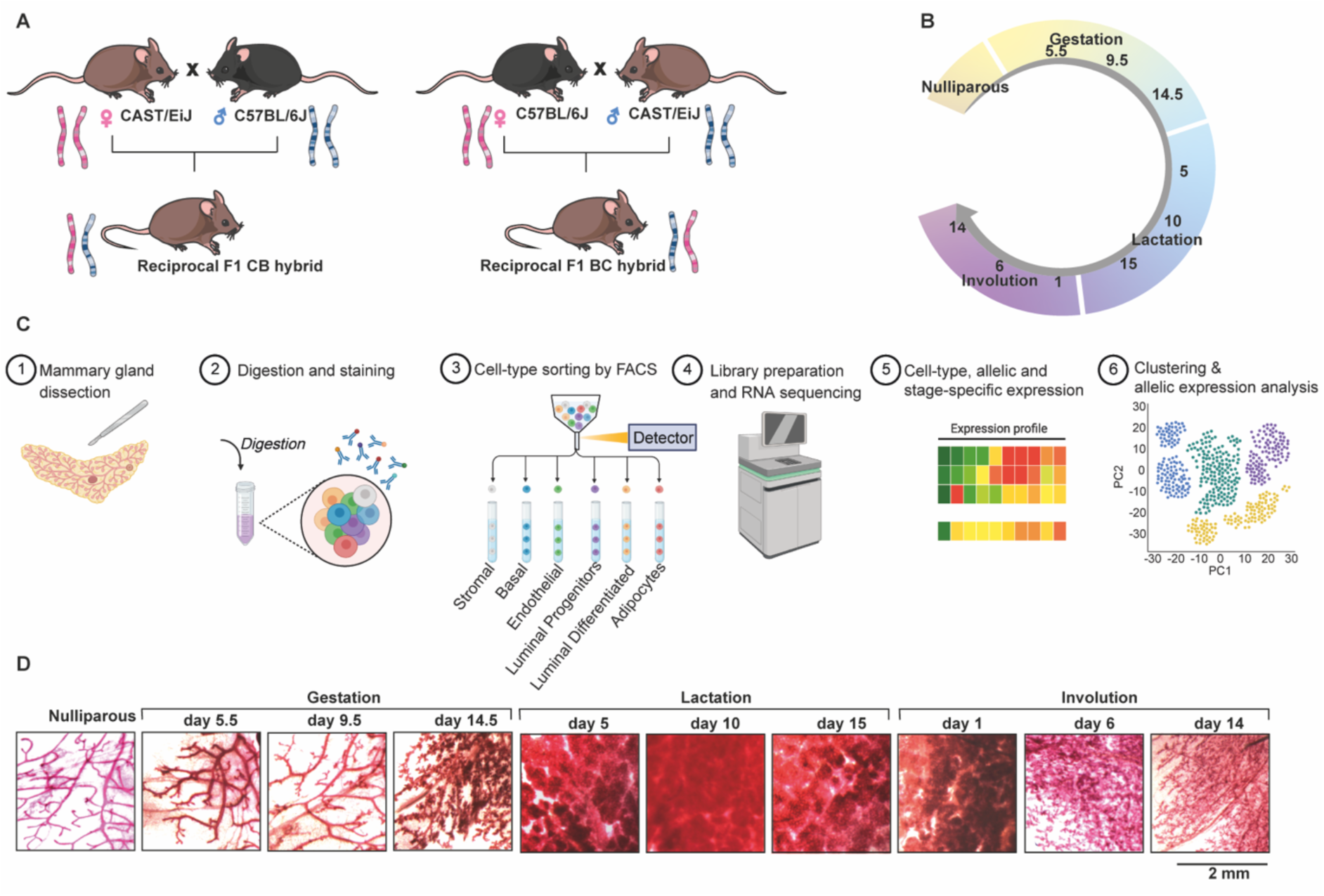
Schematic overview of the experimental strategy to generate cell-type and parent-of origin-specific expression of mammary gland cells. **(A)** Left: CAST/EiJ female mice were crossed to C57Bl6/J males to produce F1 hybrid mice, hereafter referred to as CB. Right: The reciprocal cross of C57Bl6/J females with CAST/EiJ males generated F1 CB hybrid. The first letter indicates maternal inheritance while the second denotes paternal inheritance. (**B)** Mammary glands from 80 reciprocal hybrid mice were collected at ten distinct developmental time points throughout the cyclical development of the tissue starting from the nulliparous state, at three time points during gestation (gestation day 5.5, 9.5 14.5), at three time points during lactation (lactation day 5, 10 and 15) and at three time points during involution (involution day 1, 6, 14). Overall, we collected mammary glands from 4 animals for each reciprocal cross, a total of 8 animals per time point, over the ten described time points. (**C)** Model of experimental workflow: Tissue processing steps included dissection of the tissue, followed by digestion and staining with cell-surface-specific markers as described in the methods. Single cells were then sorted using flow cytometry into specific populations, RNA was extracted and libraries for sequencing were generated. Allele-specific transcriptomes were analysed to generate and assess cell-type, stage and allele-specific expression profiles. **(D)** Carmine alum stained dissected mammary glands at the same time points showing the typical development of the tissue. Scale bar = 2 mm.

To isolate specific mammary cells, mammary glands were dissected at each time point and immediately enzyme-digested, stained and sorted based on cell surface markers for endothelial, basal, luminal progenitors, luminal differentiated and stromal cells^32^. Mammary adipocytes were processed separately and stained simultaneously for LipidTox DeepRed^33^. Total RNA Libraries were generated from sorted cell populations and sequenced. Following quality control (methods), transcriptomes were analysed for cell types, allelic and stage-specific expression (Figure 1C). The experimental design included both reciprocal hybrids, with 4 animals for each reciprocal hybrid, across 10 developmental stages and 6 sorted cell populations, resulting in a total of 480 parent-of-origin, cell-type and stage-specific transcriptomes. Histological wholemount staining confirmed the expected branching and alveolar development throughout these stages (Figure 1D, Supplementary Figure 1).

To achieve an in-depth characterisation of cell-type specific transcriptional changes, we sorted basal cells (Lin^-^CD31^neg^CD45^neg^EpCAM^lo^ CD49f^hi^), luminal differentiated cells (Lin^-^CD31^neg^CD45^neg^EpCAM^high^ CD49f ^low^ CD49b ^low^), luminal progenitor cells (Lin^-^ CD31^neg^CD45^neg^EpCAM^high^ CD49f ^low^ CD49b ^high^), immune cells (CD45^+^) and endothelial cells (CD31^+^) (Figure 2A for nulliparous stage, see Supplementary Figure 2A-C for gestation, lactation and involution). To include mammary adipocytes, representing a large fraction of mammary gland cells, we sorted simultaneously the subnatant cellular fraction (LipidTox DeepRed^+^) as previously described^33^ (Figure 2B). We next evaluated the relative percentage of each sorted cell population made up at each time point. Profiling 10 stages across the adult mammary developmental cycle revealed dynamic behaviour of the tissue, with mammary adipocytes making up the largest cell population in nulliparous and involution day 14. Immune cells show expansion during lactation, consistent with supplying immune factors from mother to offspring to support and protect them at this vulnerable stage. The luminal progenitor population shrinks in its relative proportion when transitioning from gestation to lactation, while luminal differentiated cells undergo an expansion as the tissue develops (Figure 2C). Absolute numbers of sorted cells and RNA integrity number are shown (Supplementary Table 1 and Supplementary Figure 3).

**Figure 2.**
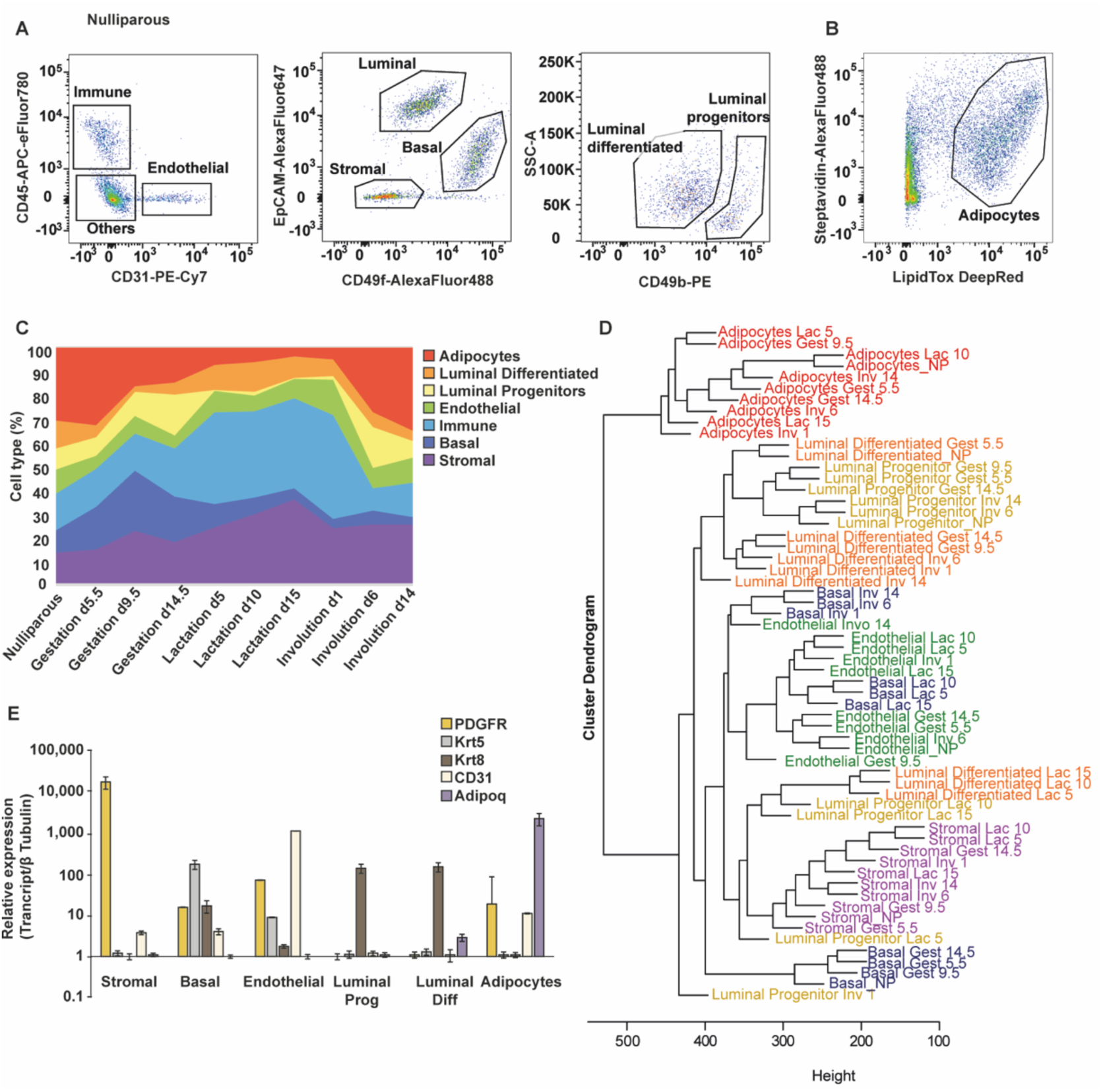
**(A)** Representative flow cytometry scatter plots for nulliparous mammary glands showing the isolated endothelial, luminal, basal and stromal cell populations. (**B)** Representative flow cytometry scatter plots for nulliparous mammary glands showing isolated mammary adipocyte population. (**C)** Dynamics of 480 cell type populations throughout the 10 developmental stages sorted in this study, represented as a percentage of total sorted cells. (**D)** Average linkage clustering of the transcriptomes from all cell types and stages performed in this study. (**E)** Expression profiles of the sorted cell populations quantified by quantitative real-time PCR in nulliparous mammary glands. n=4 biological replicates, error bars represent SEM.

To create a comprehensive atlas reflecting the dynamic transcriptional changes across the adult developmental cycle of the mammary gland we profiled the expression of all transcripts in the mouse genome. Initially, we performed average linkage clustering to identify global similarities in gene expression patterns between cell types and developmental stages. Dendrogram of natural clusters based on gene expression, revealed that most of the clustering corresponded to the cell type. Some cell types cluster distinctly based on their cellular identity rather than developmental stage, while others showed patterns influenced by both cell type and developmental stage. Transcriptionally, mammary adipocytes emerged as the most distinct cell type, clearly separating from all other cell types and stages. This suggests that mammary adipocytes undergo unique transcriptional changes throughout gestation, lactation and involution exhibiting traits different from other mammary cell types. Additionally, the stromal population clustered separately from epithelial and endothelial cells, indicating a distinct transcriptional signature likely related to their role in providing nutrition, scaffolding and supporting tissue architecture changes.

In contrast, other mammary cell types displayed more similar transcriptional landscapes. In the basal compartment, we identified clear stage-specific separation. Basal cells from the nulliparous stage clustered with those from gestation stages, but showed a substantial transcriptional shift during lactation, clustering with endothelial cells, which do not show a large transcriptional change throughout development. Involution stages again led to a unique transcriptional profile in basal cells, suggesting that their functional transition into contractile myoepithelial cells is accompanied by significant transcriptional shifts.

The luminal compartment showed closer resemblance within progenitor and differentiated luminal cells during nulliparous, gestation and involution stages, when these cell types cluster together. This indicates shared features between them, such as hormone sensing, similar to previous observations^42^. In contrast, during lactation luminal cells clustered more distantly from the non-lactating stages, supporting the notion of a major transcriptional change corresponding with secretory state and milk synthesis. Overall, lactation induced the most dramatic transcriptional changes in all cell types, affecting both epithelial and non-epithelial cells (Figure 2D).

Experimental validation of sorted cell types in nulliparous mammary glands confirmed the accuracy of our pipeline, revealing clear expression of typical markers specific to each sorted population. Specifically, the platelet derived growth factor receptor alpha (*Pdgrfa)* was highly expressed specifically in stromal cells, Keratin 5 (*Krt5)* in basal cells, Platelet and endothelial cell adhesion molecule 1(*Pecam1*/*CD31)* in endothelial cells, Keratin 8 (*Krt8)* in luminal differentiated and progenitor cells, and adiponectin (*Adipoq*) in mammary adipocytes (Figure 2E).

### Temporal expression of differentially expressed transcripts identifies potential regulators of mammary gland development

To characterise the transcriptomes, we identified all transcripts with RPKM>1 for each sorted cell population across all samples. We found a consistent number of genes detected across development and cell types, with slightly lower numbers for mammary adipocytes (Figure 3A, Supplementary Table 2). Principal component analysis (PCA) of the transcriptomes revealed clear separation by cell types and stage but not by reciprocal cross, indicating no major strain-specific differences in transcription (Figure 3B). Although well separated, luminal progenitor and differentiated cells were placed in proximity to each other, indicating a greater transcriptional resemblance between them compared to other cells.

**Figure 3.**
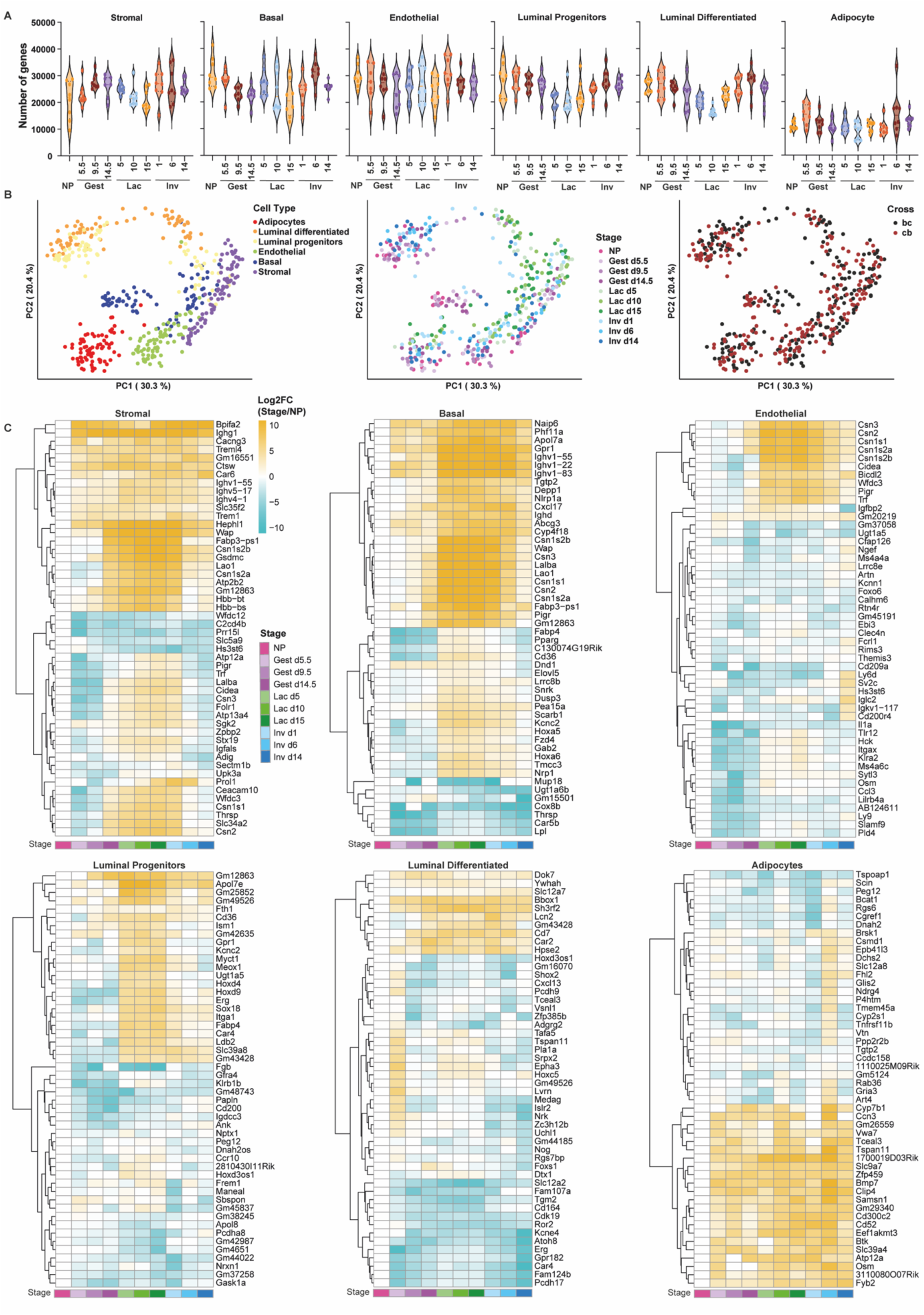
**(A)** Violin plots for the number of genes identified with RPKM>1 for each sorted cell population from 80 animals. **(B)** PCA plots of the two top principal components for all transcriptomes, colour separated by cell type (left), stage (middle) and reciprocal cross (right) **(C)** Heat maps of the top 50 most differentially expressed transcripts between stages within each cell population relative to nulliparous. Gradient scale represents log2 fold change of the time point explored relative to the nulliparous time point of the same cell type.

To explore novel regulators of mammary gland development we analysed differentially expressed genes compared to the nulliparous stage within the same cell type. This analysis generated time and cell-type-dependent progression of transcripts. We set up the top 1% of the most differentially expressed genes as a threshold and plotted the top 50 (Figure 3C). The expression pattern reveals intriguing dynamics. We chose to focus on three patterns: genes showing upregulation in gestation and peaking in lactation, genes which are robustly downregulated from gestation, and genes displaying inverse patterns of expression in different cell types.

Among the genes upregulated in gestation and peaking in lactation, we identified Whey acidic protein (*WAP*), Caseins (*Csn1, Csn2, Csn3*), Transferrin (*Trf*) and Lactalbumin (*Lalba*). These genes are known for their roles in lactogenesis and lactation. Notably, *WAP* was upregulated not only in epithelial cells but also in stromal, endothelial, and adipocyte cells. Other genes, such as L-amino acid oxidase 1 (*Lao1*), polymeric immunoglobulin receptor (*Pigr*), cell death-inducing DNA fragmentation factor, alpha subunit-like effector A (*Cidea*), and fatty acid binding protein 3, muscle and heart pseudogene 1 (*Fabp3*-*ps1*), showed similar patterns. *Lao1,* plays a role in host defence in the mammary gland^43^, *Pigr*, in transcytosis of immunoglobulins in mucosal membranes^44^, *Cidea* in lipid secretion and pup survival^45^ and *Fabp3-ps1* is a pseudogene related to fatty acid binding protein which is upregulated in lactation^46^. Additionally, we identified novel candidate genes showing the same pattern of expression, including *Gm12863*, *Gm43428*, *Gm49526*, tetraspanin 11 (*Tspan11*), WAP four-disulfide core domain 3 (*Wfdc3*), immunoglobulin heavy variable (III)-11-1 (*Ighvii-11*), ATPase, H+/K+ transporting, nongastric, alpha polypeptide (*Atp12a*), potassium voltage gated channel, Shaw-related subfamily, member 2 (*Kcnc2*)*, Cd36,* and G Protein-Coupled Receptor 1(*Gpr1/Cmklr2*).

While the functions of these genes in the mammary gland are not yet explored, some may be related to lactation. For example, *Wfdc3* is a protease inhibitor from the WAP-type four-disulfide core domain family, *Ighv-11* Immunoglobulin heavy chain gene, *Atp12a* is an ATPase responsible for K+ absorption in the colon^47^, *Kcnc2* is a potassium channel subfamily member, *Cd36* is a glycoprotein found with fatty acid binding protein in bovine milk fat globules^48^ and *Gpr1*is linked to brown adipose tissue (BAT) and glucose homeostasis^49,50^.

For genes robustly downregulated post gestation or during involution we identified *Hs3st6*, a member of the Heparan Sulfate 3-*O*-Sulfotransferase family observed in cancer but which remains poorly understood^51^ and *Peg12*, a paternally expressed gene also known as Frat3, of unknown function ^52^. This pattern of expression suggests these genes may regulate stemness or inhibit mammary differentiation or involution.

Several genes display inverse patterns of expression in different cell types, including *Osm, Erg, Car4, Fabp4, Hox3os1, Tceal3, Tgtp2, Thrsp,* and *Ugt1a5*.

*Osm* (Oncostatin M), is a member of the IL-6 cytokine family which has a role in proliferation of breast cancer cells and may play a role in post lactational regression of the mammary gland^53^. *Car4* (Carbonic Anhydrase 4), a marker for differentiation in other cell types, is also secreted in milk and regulates pH balance^54^. *Fabp4*, a circulating fatty acid-binding adipokine, is linked to mammary tumour progression^55,56^ and milk yield in dairy cows^57^. *Hox3os1*, a member of the Hox gene family, *Tceal3* (transcription elongation factor A), *Tgtp2* (T cell specific GTPase 2), *Thrsp* (Thyroid Hormone Responsive) and *Ugt1a5*, (UDP-glucuronosyltransferase) have yet to have their functions determined in the mammary gland.

The differential gene expression between cell types for each time point is provided (Supplementary Tables 3-12). Overall, this transcriptome analysis enables the identification of new candidate genes to explore in the context of lactation and mammary gland development.

### Imprinted genes are monoallelically expressed in mammary gland cells across the adult developmental cycle

We performed a comprehensive transcriptome-wide profiling of allelic expression across cell types and time points to determine the repertoire of imprinted genes in different cell types of the mouse mammary gland. We compared all known imprinted genes^58^ within 480 libraries generated from 80 animals, encompassing 6 sorted cell populations: stromal, basal, endothelial, luminal progenitors and differentiated cells and mammary adipocytes. This included two reciprocal hybrid crosses with four biological replicates per reciprocal cross. Each box represents 8 animals. To avoid lowly expressed genes, we set a threshold for gene expression levels of RPKM>1 in at least one cell type and time point.

Of the known 108 mouse imprinted genes, 75 were detected in mammary cells. Among these, 18 were paternally expressed (blue) and 10 were maternally expressed (red). Grey boxes indicate expression levels below the threshold while yellow boxes indicate biallelic expression. (Figure 4). Several imprinted genes show cell-type and stage-specific expression in the mammary gland (Supplementary Figure 4).

**Figure 4.**
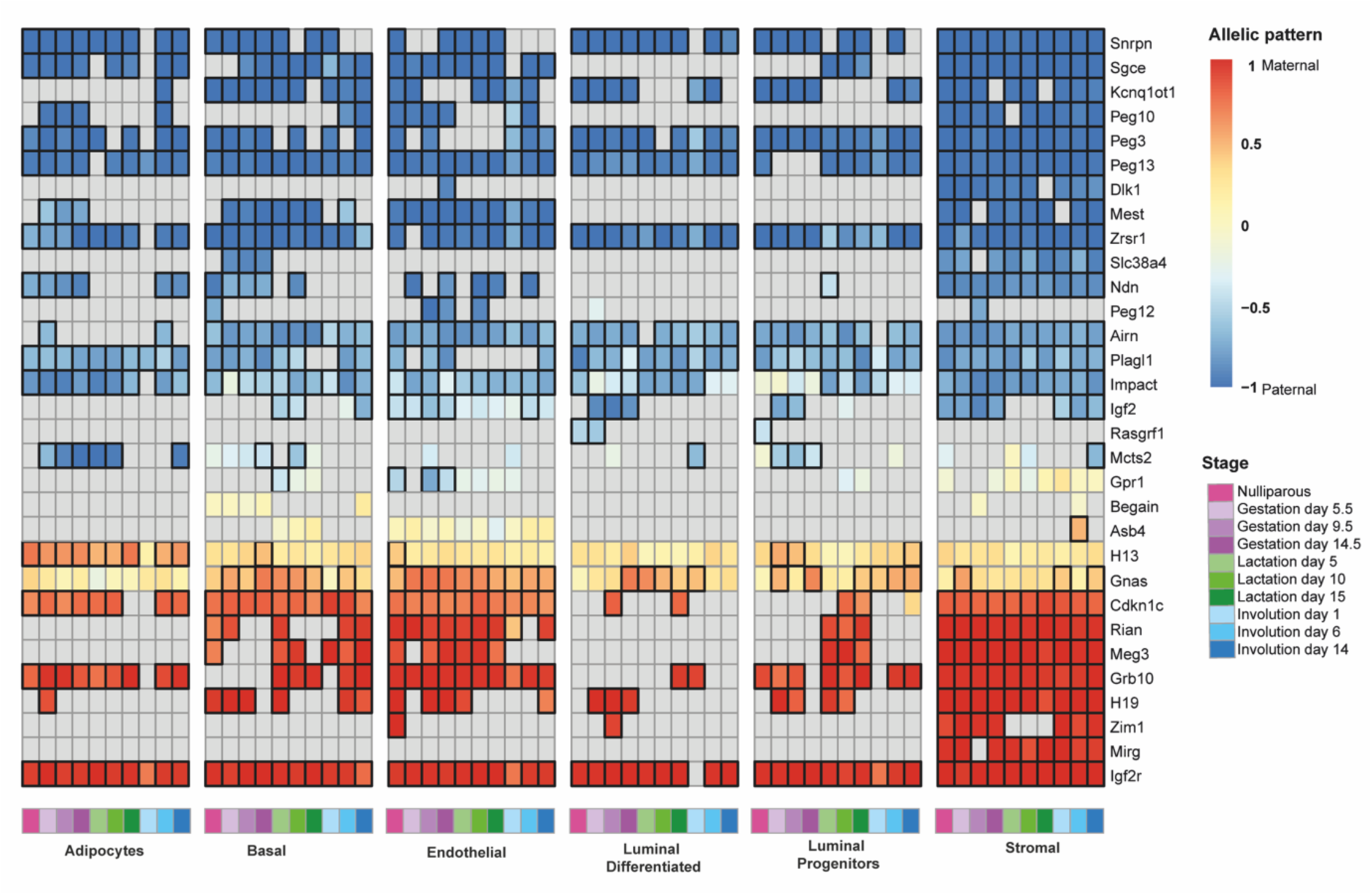
Heat map of allelic bias for all known imprinted genes which are expressed above a threshold of RPKM>1 in at least one cell type across the adult mammary gland developmental cycle. Maternal and paternal expression are indicated in red and blue respectively, while yellow indicates biallelic expression. Grey indicated an expression level lower than the threshold. Boxed cells highlight values exceeding the 0.2 or 0.7 threshold for monoallelic expression, and each box represents an average of 8 biological replicates, 4 of each reciprocal cross.

Canonical imprinted gene pairs, including *H19*-*Igf2*, *Igf2r*-*Airn*, *Meg3*-*Dlk1*, *Cdkn1c*-*Kcnq1ot1* and *Zim1*-*Peg3*, exhibited parent-of-origin-specific expression consistent with their established patterns (Supplementary Figure 5).

Several imprinted genes are monoallelically expressed in specific mammary cells throughout development, suggesting potential roles in regulating mammary gland development, lactogenesis or milk composition. Examples include *Igf2r*, *Grb10*, *Meg3*, *Rian*, *Cdkn1c*, *Mest*, and *Sgce*. Interestingly, some of these genes show robust allelic expression in luminal or basal lineages which play major roles in milk synthesis and let-down.

The patterns of gene imprinting and allele-specific expression in the mammary gland present a nuanced picture of transcriptional regulation: For example, *Gnas*, a transcript-specific imprinted gene, shows high maternal expression in endothelial, basal and luminal cells, but in mammary adipocytes, expression displayed only a maternal bias. This pattern is consistent with previous observations in other cell types, where *Gnas* is preferentially or exclusively expressed from the maternally inherited chromosome^59^.

*Asb4*, previously reported as imprinted in the embryo^60^ and adult^61^, exhibited high expression in endothelial and basal cells, but showed biallelic expression in the mammary gland, unlike its canonical imprinting pattern. *Begain*, a gene known to display transcript-specific paternal expression^62^, expresses the biallelic non-imprinted transcript in the mammary gland with high expression in basal cells.

The interplay between the retrotransposed *Mcts2* and its host gene *H13* provides further insights into cell-type specific monoallelic expression. While *Mcts2* is uniformly expressed across cell types, it exhibits clear paternal expression only in adipocytes and luminal progenitors during gestation. In contrast, *H13* shows consistently high expression in mammary cells throughout the developmental cycle (Supplementary Figure 4), with maternal monoallelic expression occurring specifically in adipocytes and a maternal bias elsewhere. *Gpr1*, typically biallelic in most tissues and reported as paternally-expressed explicitly in the kidney^63^, showed paternal expression in mammary endothelial cells, where it is highly expressed particularly during lactation.

We compared imprinted gene expression in the mammary gland to their behaviour in other tissues. Of 24 genes previously identified as monoallelically expressed in the placenta, most showed either biallelic expression or weak allelic bias in mammary cells. Similarly, 5 brain-specific imprinted genes exhibited biallelic expression in the mammary gland, consistent with previously reported tissue-specific imprinting. Similarly, stage-specific or weakly biased imprints also showed biallelic expression in this tissue, (Supplementary Figure 6).

No novel imprinted genes were consistently detected across all cell types (Supplementary Figure 7), however *B830012L14Rik*, a transcript located between *Rian* and *Mirg*, is maternally expressed, likely representing another contiguous transcript from the Dlk1-Dio3 domain rather than an independently regulated gene. Previously characterised novel imprinted genes which are singletons^58^ are either not expressed or not monoallelically expressed in mammary cell types.

Imprinted gene expression in all cells of the tissue, would imply a tissue-specific function which is unique to a particular imprinted gene, while cell-type specific function would suggest a more specialised role within cellular compartments. Our data shows that most imprinted genes are expressed in specific cell types, without changes in the extent of their biased expression. This suggests that in the mammary gland, most imprinted genes are regulated at the transcriptional level and their imprinting status remains constant.

Comparison to previous work^24^ reveals several additional imprinted genes showing parent-of-origin specific expression. In this study, we identified 18 imprinted genes expressed from the paternally inherited chromosome, and 7 maternally expressed imprinted genes. Among these, three paternally expressed genes had not been detected in other studies^24^: *Dlk1*, *Peg12* and *Mcts2*. The vertebrate specific atypical Notch ligand *Dlk1*, was expressed almost exclusively in stromal cells, with very low levels in other mammary gland cells. *Peg12* displays its highest expression in endothelial cells, with moderate expression in basal and stromal cells, as well as at several points in luminal progenitors. *Mcts2*, shows paternal allelic expression in mammary adipocytes.

The cell-type expression of several imprinted genes in the mammary gland suggests that they may be tightly regulated in the mammary gland to important roles in this tissue. The consistent imprinting status maintained across different cells within the same tissue implies a stable imprint regulatory mechanism, potentially providing robust control over key aspects of mammary gland physiology. This opens avenues for further research into the specific roles of these imprinted genes in this tissue.

### Validation of parent-of-origin expression bias and identification of strain-specific expression variations

To validate allelic expression in sorted mammary gland cell types, we selected 6 imprinted genes, 3 maternally and 3 paternally expressed, and examined their allelic expression across mammary gland development by allele-specific qPCR. (Figure 5A). Pyrosequencing validation in sorted basal and endothelial cells confirmed expression from the maternally inherited chromosome of *Cdkn1c*, *Meg3* and *H19*. Conversely, *Dlk1*, *Igf2* and *Snrpn* exhibited clear paternal expression in stromal and endothelial cells (Figure 5B).

**Figure 5.**
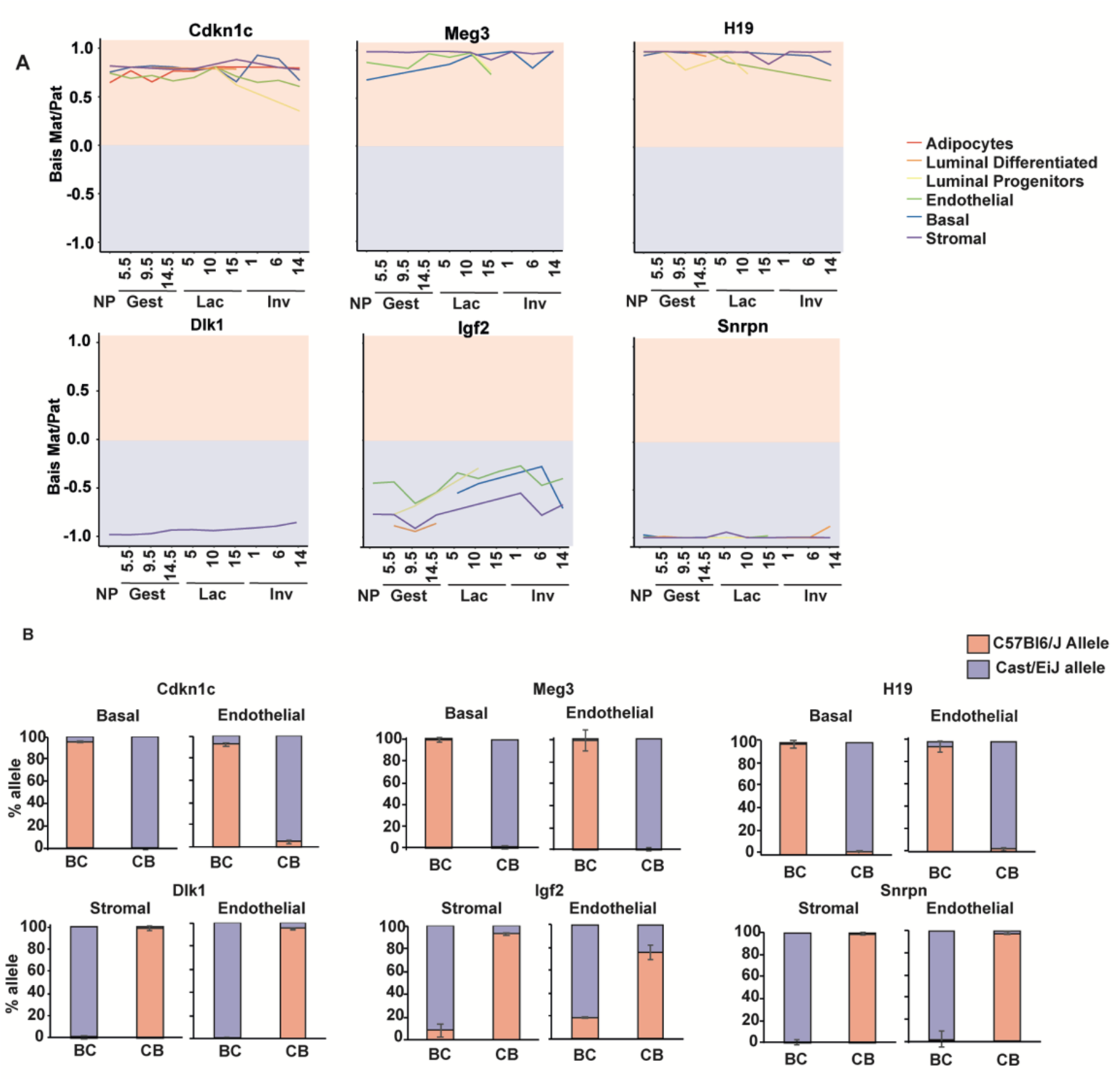
**(A)** Allelic bias dynamics of selected imprinted genes across mammary gland development and cell types. **(B)** Pyrosequencing validation of selected imprinted genes in sorted nulliparous mammary cell populations.

To assess the functional impact of genetic diversity in C57BL/6J and CAST/EiJ mice, we sought to identify genes with strain-specific expression bias and differentiate them from potential novel imprinted genes. We focused on genes with RPKM >1 in at least three time points or cell types that also displayed an expression bias greater than 0.5 to obtain the transcripts with the greatest strain-specific allelic bias. Our analysis identified 16,432 genes (Supplementary Figure 8A) potentially regulated by strain-specific enhancers or post-transcriptional mechanisms unique to one of the mouse strains indicating extensive strain-specific effects on transcript profiles.

Several strain-specific transcriptional profiles exhibit cell-type-specific allelic bias, suggesting localised and selective regulation that may vary between strains. These observed differences could also arise from variations in expression levels affecting our analysis. While previous studies_24,64_ have documented allelic bias on a tissue-specific level, our findings highlight that cell-type-specific factors within the same tissue can also influence allelic bias. A comparative analysis of strain-specific bias reveals a notable increase in the number of genes identified in our study versus previous work. Specifically, our study identified 16,432 genes with strain-specific bias, compared to 1,142 genes reported by Andergassen et al. Of these, 826 genes were common between the two studies (Supplementary Figure 8B). Additionally, we identified 2476 genes with parent of origin-specific bias, compared to 365 reported by Edwards et al., and 77 of them were common (Supplementary Figure 8C). This significant difference underscores the enhanced resolution and depth of our analysis in uncovering strain-specific expression patterns.

### Expression patterns of genes linked to lactation disorders

Mammary gland development, critical for lactation, involves alveologenesis and lactogenesis. Impaired lactation, affecting 1 in 20 women, is characterised by delayed lactogenesis or insufficient milk production, often influenced by maternal conditions such as obesity, diabetes, and thyroid dysfunction^65^. Genome-wide association studies (GWAS) have identified genetic factors underlying breastfeeding traits, including SNPs in *OXE1* (Oxoeicosanoid Receptor 1) gene, a receptor for polyunsaturated fatty acids found in breastmilk, and genes in the fatty acid desaturase family (*FADS1, FADS2, FADS3*) which influence breastmilk fatty acid composition and breastfeeding duration^66,67^.

To explore genetic links to lactation, we integrated GWAS findings with cell-type-specific mammary gland transcriptomes across ten developmental stages. Using the NHGRI-EBI catalogue, we identified SNPs associated with the traits “breastmilk measurements” and “breastfeeding duration” and analysed the expression patterns of these SNP-hosting genes. Our analysis identified several differentially expressed genes in mammary epithelial cells, particularly luminal cells, during lactation. These genes are involved in the cellular metabolism, differentiation and structural remodelling, critical for milk production (Figure 6A).

**Figure 6.**
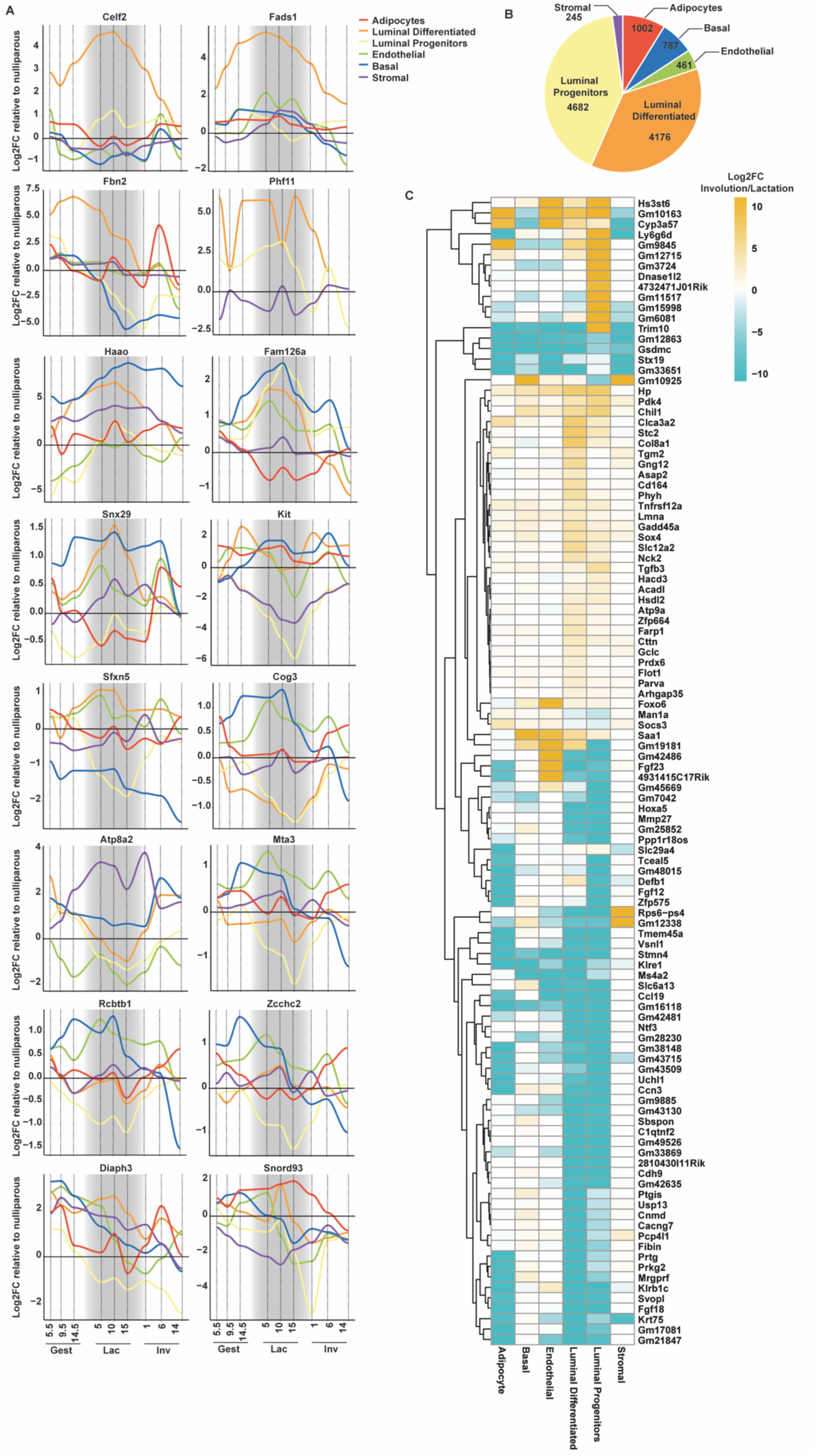
**(A)** Expression pattern of differentially expressed genes (DEGs) containing SNPs associated with the traits “breastmilk measurements” and “breastfeeding duration”, relative to nulliparous state, across mammary gland developmental stages and cell types. The lactation phase is highlighted with grey shading. **(B)** Numbers of DEGs identified per cell type containing SNPs associated with breast cancer. **(C)** Heat maps of the most differentially expressed genes (Padj <=10^-17^) comparing lactation and involution across cell populations containing SNPs linked to breast cancer.

#### Upregulated Genes in Luminal Cells During Lactation

Genes supporting milk production were highly expressed in luminal cells during lactation. *Celf2*, an RNA-binding protein, facilitates alternative splicing enabling rapid protein synthesis ^68^. *Fads1*, critical for long-chain polyunsaturated fatty acid (LC-PUFA) synthesis, enhances milk fatty acid content ^69^. Its knockout in mice results in early mortality, reversible by arachidonic acid supplementation ^70^. *Fbn2*, encoding fibrillin-2, supports extracellular matrix elasticity ^71^, which is likely essential for milk ejection. Additional upregulated genes include *Phf11*, a transcriptional co-activator of cytokines^71^, *Haao*, supporting the energetic demands of milk production in mammary cells^72^, and *Fam126a* and *Snx29* both involved in membrane trafficking and cell signalling ^73^.

#### Downregulated Genes in Luminal Progenitor Cells During Lactation

Several of the GWAS-associated genes exhibited reduced expression particularly in luminal progenitor cells during lactation, reflecting the shift from cellular proliferation and differentiation to milk synthesis. *Kit*, a marker of luminal progenitors^74^ was downregulated, indicating differentiation into mature luminal cells and reduced progenitor activity to minimise abnormal growth. Decreased expression of *Sfxn5*, involved in mitochondrial homeostasis^75^, *Cog3*, a regulator of Golgi vesicle trafficking ^76^, *Atp8a2*, which maintains membrane asymmetry^77^, *Mta3*, involved in epithelial-mesenchymal transition (EMT) ^78^ and *Rcbtb1*, a ubiquitin ligase regulator^79^, aligns with the terminal differentiation of luminal cells. This shift corresponds to a reduction in developmental EMT pathways during lactation.

#### Downregulated Genes During Mammary Involution

During mammary gland involution, genes associated with structural remodelling and cellular processes displayed decreased expression. For example, *Diaph3*, involved in actin cytoskeleton remodelling^80^, was significantly downregulated in luminal cells, reflecting reduced cytoskeletal activity during apoptosis. However, its upregulation in mammary adipocytes suggests a role in niche repopulation. Moreover, *Snord93*, a gene involved in RNA modification, was downregulated during involution, consistent with a potential role in RNA degradation during apoptosis.

#### Predictive Gene Signatures for Postpartum Breast Cancer

Lactation transforms the mammary gland, involving significant morphological, transcriptional, and epigenetic changes^81,82^. Breastfeeding benefits maternal health by reducing breast cancer risk, though the mechanisms behind this protection remain unclear^83^. Research has shown that shorter breastfeeding durations are linked to more aggressive breast cancer subtypes, such as triple-negative and basal-like, with poorer prognoses^84,85^. Postpartum breast cancers (PPBC), occurring within 5–10 years of birth, are linked to hormonal fluctuations, tissue remodelling, and pro-inflammatory microenvironments during mammary gland involution, making the gland more susceptible to malignancy^86^.

Mammary involution, marked by cell clearance and extracellular matrix remodelling, creates conditions that may foster tumour progression. Milk stasis exacerbates inflammation, which, alongside immune suppression, contributes to the development of PPBC ^87^. Luminal epithelial cells and mammary adipocytes are implicated in these processes, but the molecular and genetic mechanisms remain poorly understood^88–90^.

To investigate PPBC gene signatures, we integrated GWAS data with postpartum mammary gland transcriptomes. SNPs associated with breast cancer were mapped to 21,975 protein-coding transcripts, revealing 11,353 differentially expressed genes (DEGs) during the transition from lactation (day 10) to involution (days 1, 6, and 14; Padj < 0.05). Luminal cells exhibited the highest number of DEGs, consistent with their central role in breast cancer development. Mammary adipocytes also showed significant SNP-linked DEGs, underscoring their emerging relevance in PPBC 91 (Figure 6B).

Focusing on the most significant changes (Padj<10^-18^), we identified 76 genes potentially linked to PPBC (Figure 6C). Many were downregulated in luminal cells during involution, likely reflecting the transcriptional burst in lactation followed by apoptosis during weaning. For instance, *MMP27*, while not directly linked to breast cancer, belongs to the matrix metalloproteinase family associated with tumour progression. Similarly, *Hoxa5*, upregulated in luminal cells during involution, has been associated with ER+ breast cancer^92^.

Upregulated genes included *FoxO6* in endothelial cells, a transcription factor with limited evidence linking it to breast cancer^93^, and *Hs3st6*, encoding an enzyme involved in heparan sulfate biosynthesis, which may influence tumour cell proliferation. While related enzymes promote breast cancer cell proliferation^94^. *Gsdmc*, linked to other cancers ^95^, and *Stx19*, a gene with little prior breast cancer research ^96^, also warrant further investigation.

In summary, our analysis provided insight into the dynamics and specificity of mouse genes with both established and novel associations with breast cancer in their human counterparts. While some genes have known functions in malignancy, others remain unexplored, offering valuable targets for future studies on the mechanisms underlying PPBC.

## Discussion

This study presents a comprehensive atlas of imprinted genes in the mammary gland, establishing a foundation for further exploration of their roles in lactation and postnatal development. Utilising extensive transcriptome data, we provide temporal expression profiles of known and novel candidate genes that may regulate mammary gland development. Additionally, we identify and validate imprinted genes, characterising the mouse mammary gland allelome throughout the adult mammary developmental cycle, a crucial component previously missing in other studies^24,25,42,64^.

Our flow cytometry analysis sheds light on the cellular dynamics within the tissue, with a particular focus on understudied mammary non-epithelial cells. Despite the novel transcriptome data generated from mammary adipocytes, several technical challenges impact our findings in that cell-type. We isolated mammary adipocytes using a lipid dye, providing a general overview of imprinting and transcriptional dynamics in these cells, which form a significant part of the gland at various stages. However, the lack of cell-type-specific surface markers for flow cytometry limits our ability to separate sub-populations of mammary adipocytes or identify developmental changes in their identity. The technical challenges in sorting mammary adipocytes result in a slightly reduced yield, and their lower RNA content affects the quality of our libraries, leading to fewer detected genes. Additionally, immune cells identified as CD45^+^ through flow cytometry were not sequenced due to their heterogeneous nature and the risk of mixing tissue-resident and non-resident immune cells, particularly during lactation.

We observed a robust pattern of parent-of-origin expression and the influence of cell type on allelic bias. Analysis indicates that most changes in gene expression occur between the nulliparous and lactation stages, with gestation and involution often displaying similar expression patterns. Notably, we identified several additional imprinted genes in the mammary gland compared to previous studies^24^, attributed to the higher cellular resolution of this study. At the same time, we profiled and expanded the knowledge on strain-specific expression bias. This study, therefore, provides an unprecedented level of detail and sets the stage for uncovering the physiological significance of specific imprinted genes in the mammary gland.

While our transcriptome data is deep, some imprinted genes show dynamic expression in vivo during pregnancy and lactation, with low expression levels making it difficult to determine allele-specific expression in the analysis. Certain instances remain challenging for determining if a gene is imprinted due to specific splice variants, such as in the case of *Gnas*. Long-read sequencing would better address these splicing issues, providing more precise insights.

Our validation using pyrosequencing reveals a higher percentage of allelic bias in most tested genes, indicating that our allele-specific transcriptome may underestimate allelic expression. Pyrosequencing, being more sensitive, detected *Dlk1* not only in stromal cells, as seen in the transcriptome, but also in endothelial cells that did not meet the analysis threshold. Similarly, *Igf2* showed stronger pattern-of-origin expression in pyrosequencing compared to RNA sequencing.

The prevailing theory about the evolution of genomic imprinting suggests that imprinting is driven by a conflict between maternal and paternal genes over resource allocation to offspring^10,11,13^. This hypothesis has been developed based on a relatively small subset of genes and focused on tissues influencing offspring growth such as the placenta, a site for potential parental conflict, where some paternally expressed genes promote growth and resource acquisition, while some maternally expressed genes act to suppress resources allocation, to preserve energy for future offspring.

However, the mammary gland represents a unique case where this conflict does not follow this pattern, as the tissue retains grandpaternal expression of paternal genes, less directly related to the paternal genome of the conceptus. This may indicate a different regulatory mechanism and challenge the generality of the conflict hypothesis. It may be that imprinting dynamics in the mammary gland evolved under different selective pressures, related to its role in nurturing and influencing the health and development of future progeny.

Results suggest that the selective absence of imprinting is a mechanism to control gene dosage specifically in the brain rather than a general phenomenon maintaining stemness. This finding aligns with previous studies^64^, which indicate that tissue-specific imprinting is rare, with most genes being imprinted across all adult tissues where they are expressed. Previous work has highlighted selective absence of imprinting in specific tissues, such as *Igf2* in the neural stem cells of the hippocampus, where, in contrast to the choroid plexus and leptomeninges, it maintains its imprinting when acting as an autocrine/paracrine factor with an essential mitogenic role in neurogenesis. However, in our study, we observed an increase in *Igf2* dosage in luminal cells without detecting any change in paternal expression, suggesting that the selective absence of imprinting does not act in the mammary gland.

Stroganstev et al.^97^ found strain-specific variation in the binding motif of ZFP57 between CAST/EiJ and C57BL/6J mice, with genetic variation outside the motif potentially determining the methylation state and subsequently influencing strain-specific gene expression bias. Consistent with this, we observed a greater strain bias towards the CAST/EiJ allele compared to the C57BL/6J allele, which may reflect an alignment bias in our data.

By integrating GWAS data with mammary gland-specific transcriptomes, we identified expression patterns of genes linked to lactation disorders, and predictive gene signatures for PPBC. We identified key genes that are up or downregulated, revealing potential pathways that may be critical for milk synthesis, secretion, and structural remodelling. Furthermore, we explore the links between lactation, involution and the increased risk of PPBC, and identify DEGs during the lactation-to-involution transition, with a notable enrichment in luminal epithelial cells and adipocytes. The number of SNPs available to us was significantly higher for breast cancer compared to lactation disorders. This reflects a bias in research of breast cancer compared to normal mammary gland biology and specifically lactation and may impact our analysis. Nevertheless, this could inform future research regarding candidate genes specifically involved in PPBC and lactation disorders. Future studies are required to understand the functions of the identified genes to inform public health strategies and personalised lactation support.

This dataset is a valuable resource for future studies on transcriptional allelic expression and dynamics in different cell types in the mammary gland. It highlights imprinted genes that are monoallelically expressed in mammary cells and can regulate postnatal nutritional resources and allows identification of candidate imprinted genes for further exploration such as *Cdkn1c*, a cyclin-dependent kinase inhibitor that regulates the relationship between proliferation and differentiation, and *Dlk1*, which can maintain cells in quiescence and negatively regulate differentiation. Our results provide a global, unbiased view of adult mammary gland development and extend to the non-imprinted transcriptome, offering new insights into nutrient transfer between mother and offspring.

---

**Supplementary Table 1**

Absolute sorted cell numbers from each animal used in this study, including data on the reciprocal cross and stage.

**Supplementary Table 2**

The absolute number of genes with RPKM>1 separated per animal and cell type and of them, the number of genes containing a SNP which were used subsequently for the allele-specific expression analysis.

**Supplementary Table 3-12**

Differentially expressed genes between cell types within each stage.

**Supplementary Table 13**

Primer sequences used in this study.

## Author contributions

ACFS and GH planned the experiments and interpreted the data. GH performed all the experiments, animal work and sample processing, analysed the data and wrote the manuscript. GH and KRC analysed the data and prepared the figures. All authors contributed to the writing and editing of the manuscript. KRC and HT performed RNA-seq expression analysis and allelic-expression analysis. BA explored GWAS datasets, CAE contributed to the analysis of imprinted genes, SP assisted with RNA processing and library preparations.

## Data Availability

RNA-Seq data were deposited into the NCBI BioProject under accession number PRJNA1231051. All other data supporting the findings of this study are available within the paper and its Supplementary Information.

## Acknowledgements

This work was funded by MRC grants MR/R009791/1 to ACF-S and MR/W003783/1 to ACF-S and GH. G Hanin was supported by the Royal Society Newton Fellowship and FEBS long-term fellowship.

The Cambridge NIHR BRC Cell Phenotyping Hub, and the Cambridge Combined Animal Facility supported the project.

## Supplementary data

**Supplementary Figure 1.**
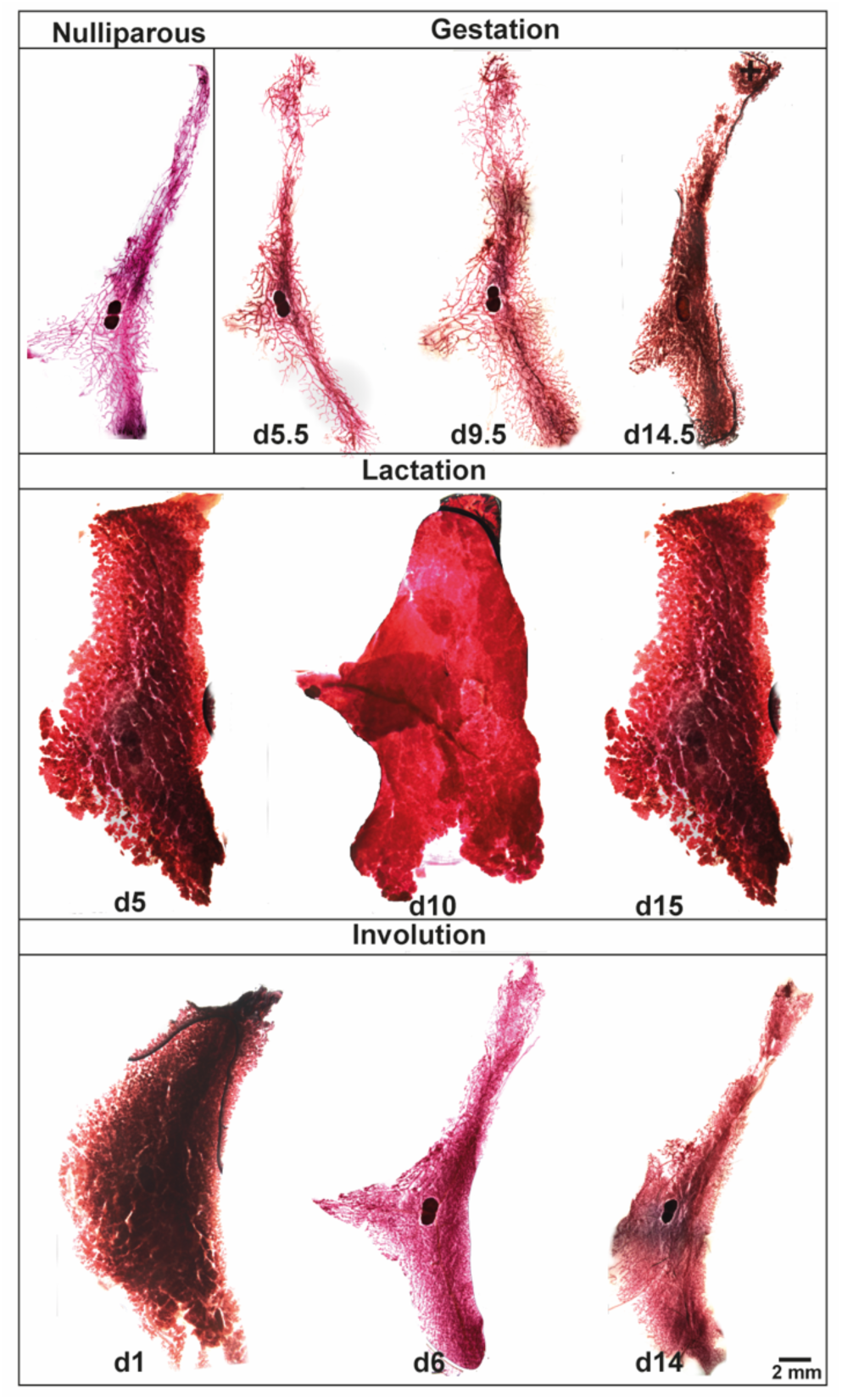
Carmine alum stained wholemounts of dissected mammary glands at the same time points showing the typical development of the tissue, showing entire images of abdominal mammary glands.

**Supplementary Figure 2.**
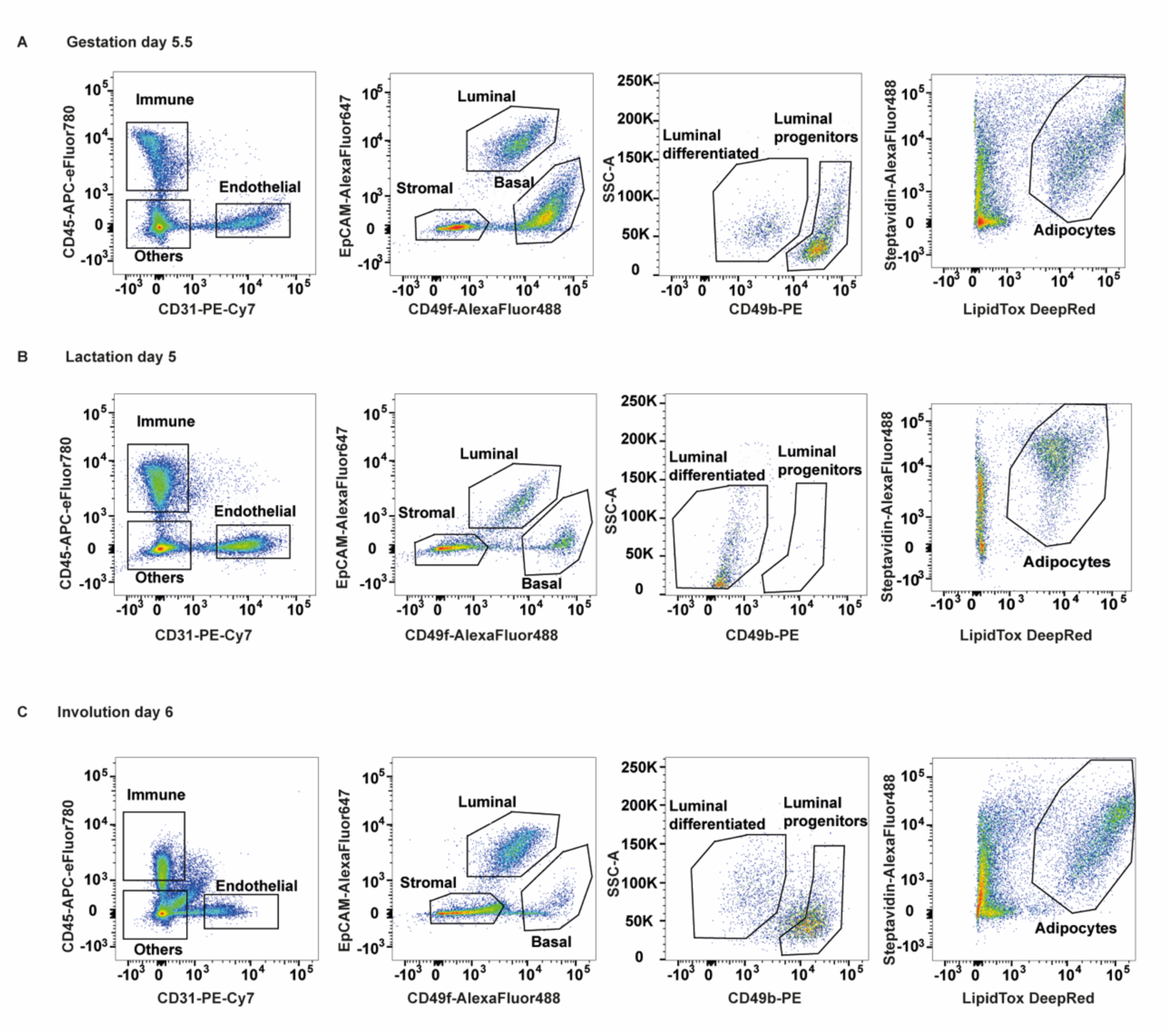
A-C Representative flow cytometry dot plot for gestation day 5.5 **(A)**, lactation day 5 **(B)**, and involution day 6 **(C)** mammary glands showing the isolated endothelial, luminal, basal, stromal, and adipocyte cell populations.

**Supplementary Figure 3.**
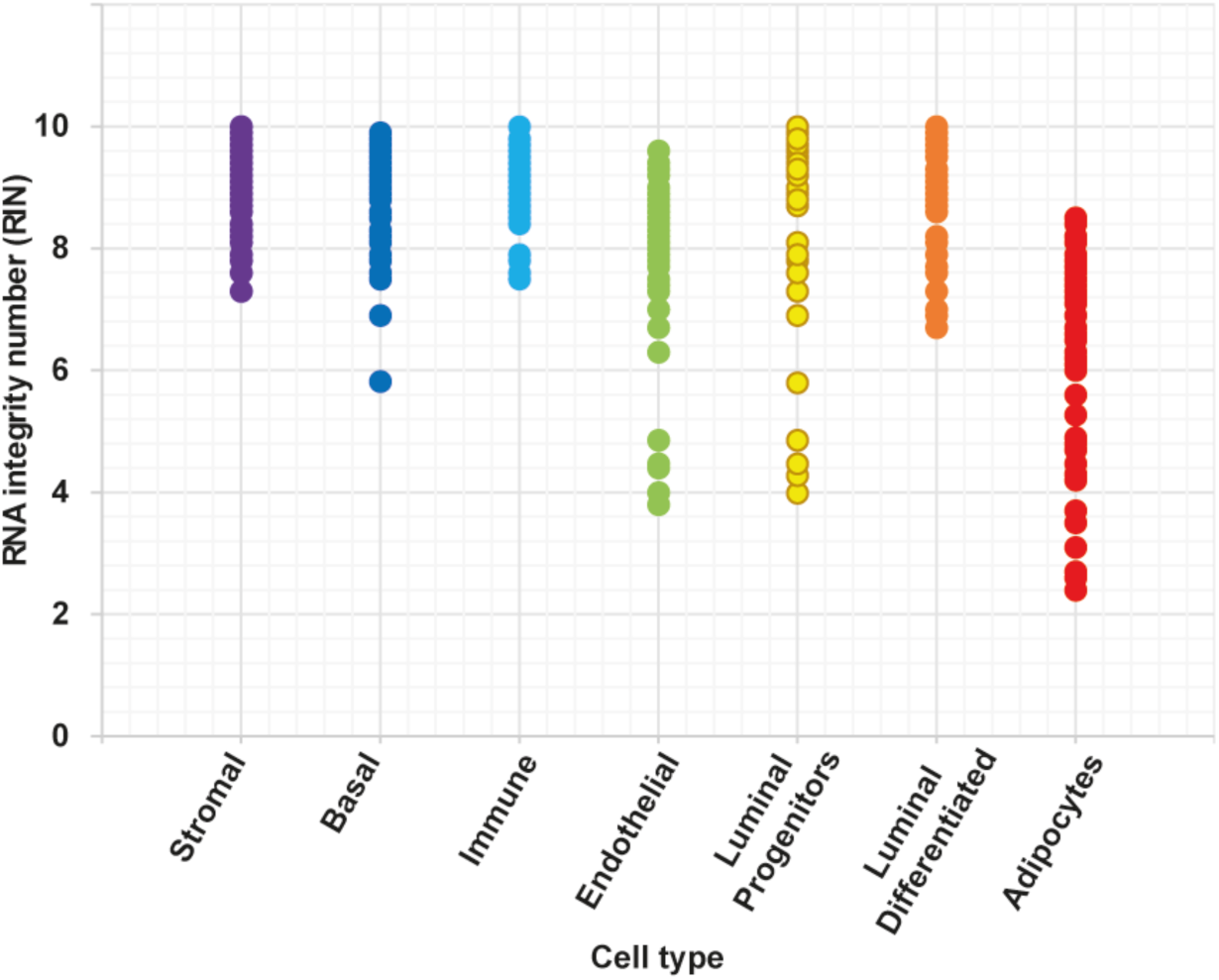
RNA integrity number of 80 sorted cell populations used in this study, including stromal, basal, immune, endothelial, luminal progenitor and differentiated cells and mammary adipocytes. Each dot represents the RNA integrity number of a population used for RNA-seq library preparation.

**Supplementary Figure 4.**
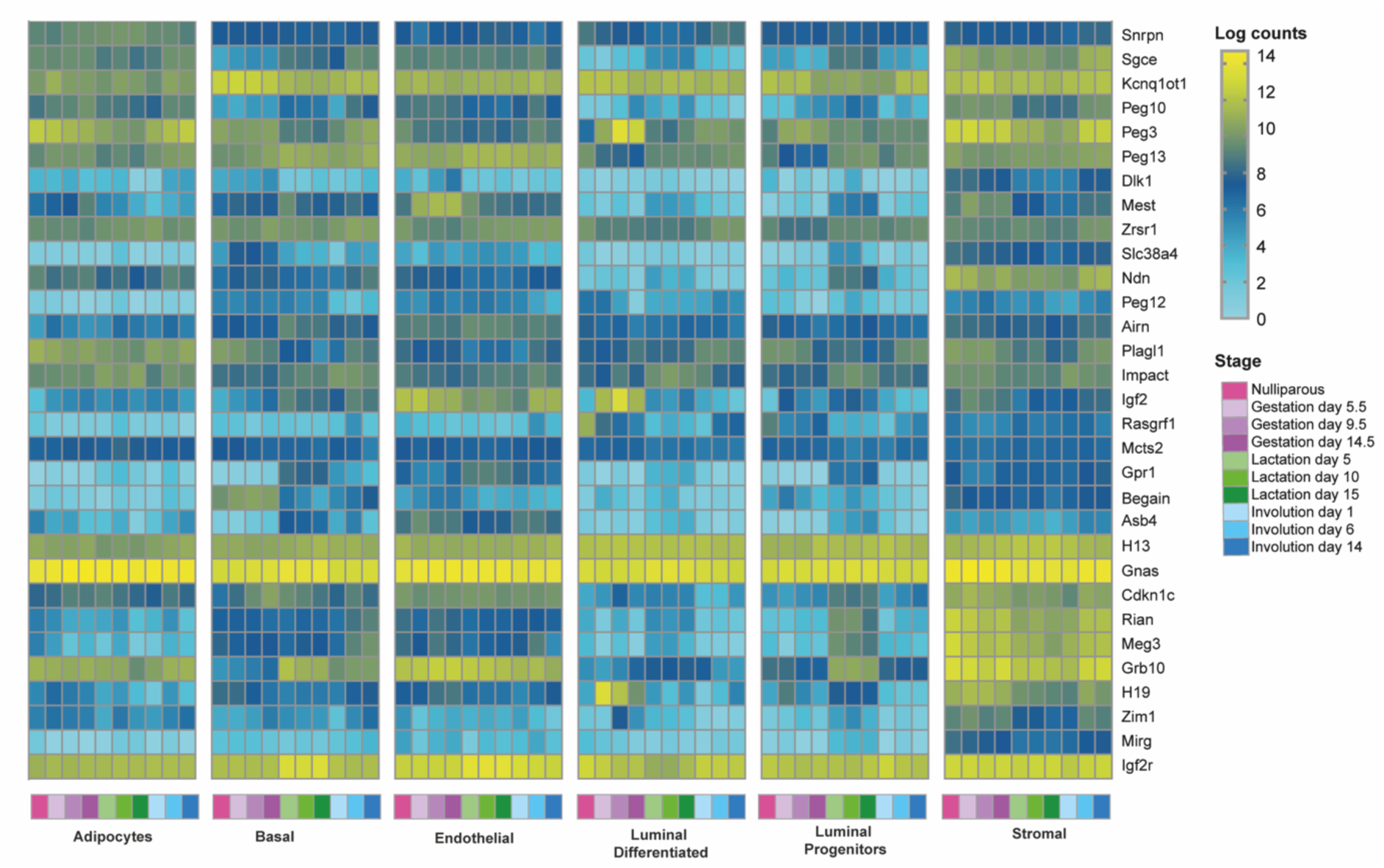
Expression levels of imprinted genes with allelic expression in mammary gland cells are shown across the adult mammary gland developmental cycle. Expression is represented as log transformed counts, with yellow indicating high expression level. Each box represents an average of 8 biological replicates, 4 of each reciprocal cross.

**Supplementary Figure 5.**
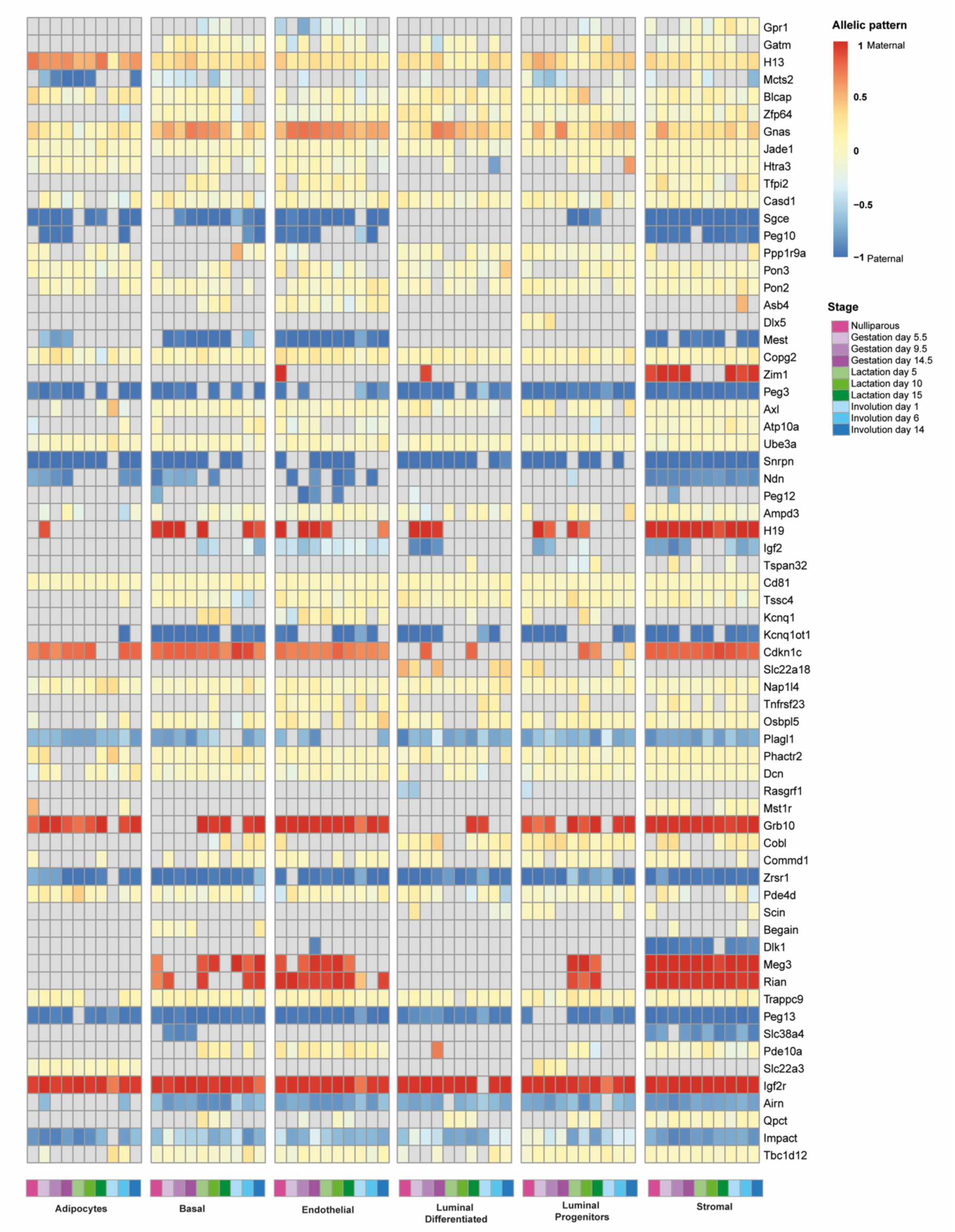
Heat map of allelic pattern arranged by chromosomal location, for all known imprinted genes which are expressed above a threshold of RPKM>1 in at least one cell type across the adult mammary gland developmental cycle. Maternal and paternal expression are indicated in red and blue respectively, while yellow indicates biallelic expression. Grey indicated an expression level lower than the threshold. Each box represents an average of 8 biological replicates, 4 of each reciprocal cross.

**Supplementary Figure 6.**
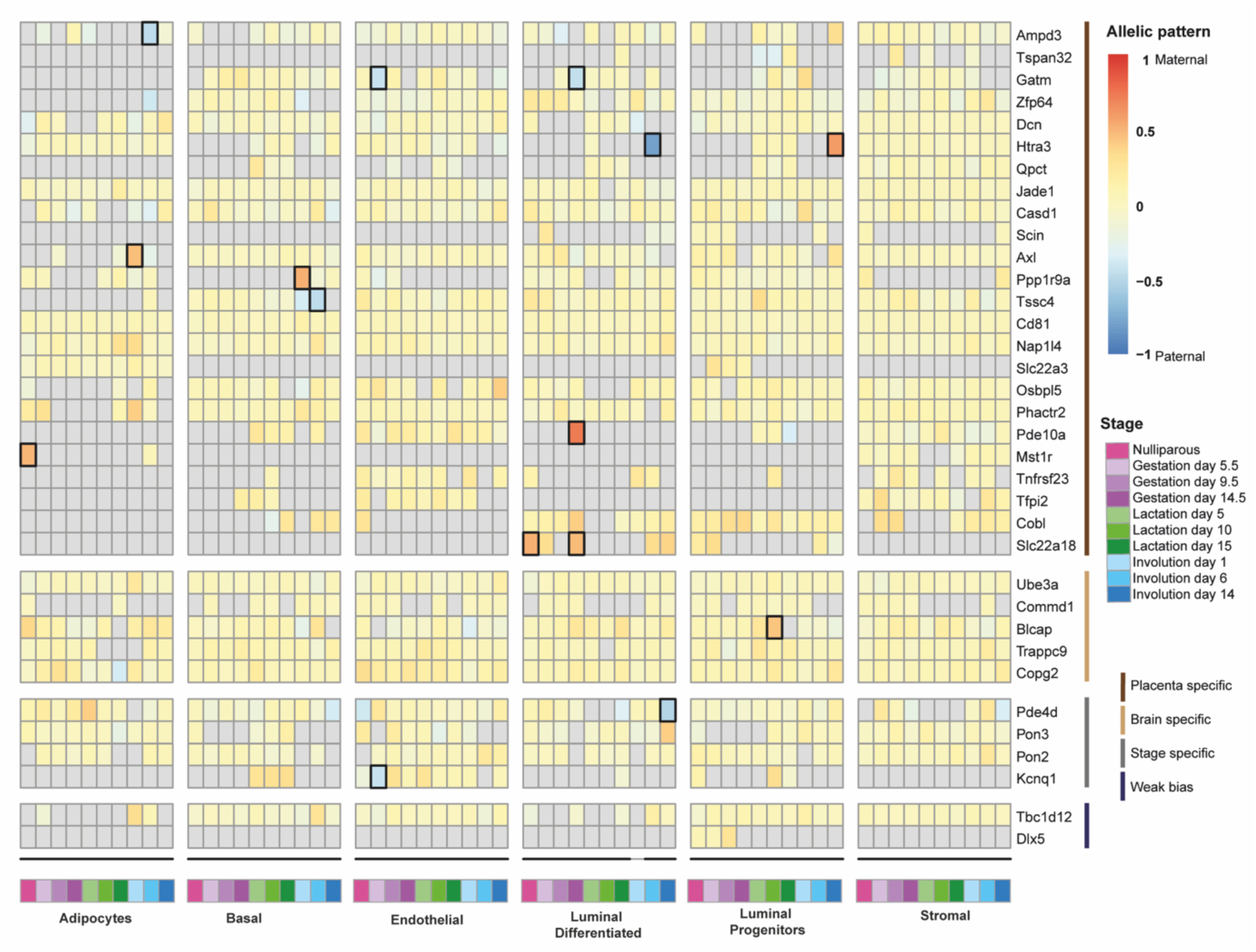
Heat map of allelic bias for imprinted genes known as placental, brain-specific, stage specific or displaying weak-bias, expressed above a threshold of RPKM>1 in at least one cell type across the adult mammary gland developmental cycle. Maternal and paternal expression are indicated in red and blue respectively, while yellow indicates biallelic expression. Grey indicated an expression level lower than the threshold. Boxed cells highlight values exceeding the 0.2 or 0.7 threshold for monoallelic expression, and each box represents an average of 8 biological replicates, 4 of each reciprocal cross.

**Supplementary Figure 7.**
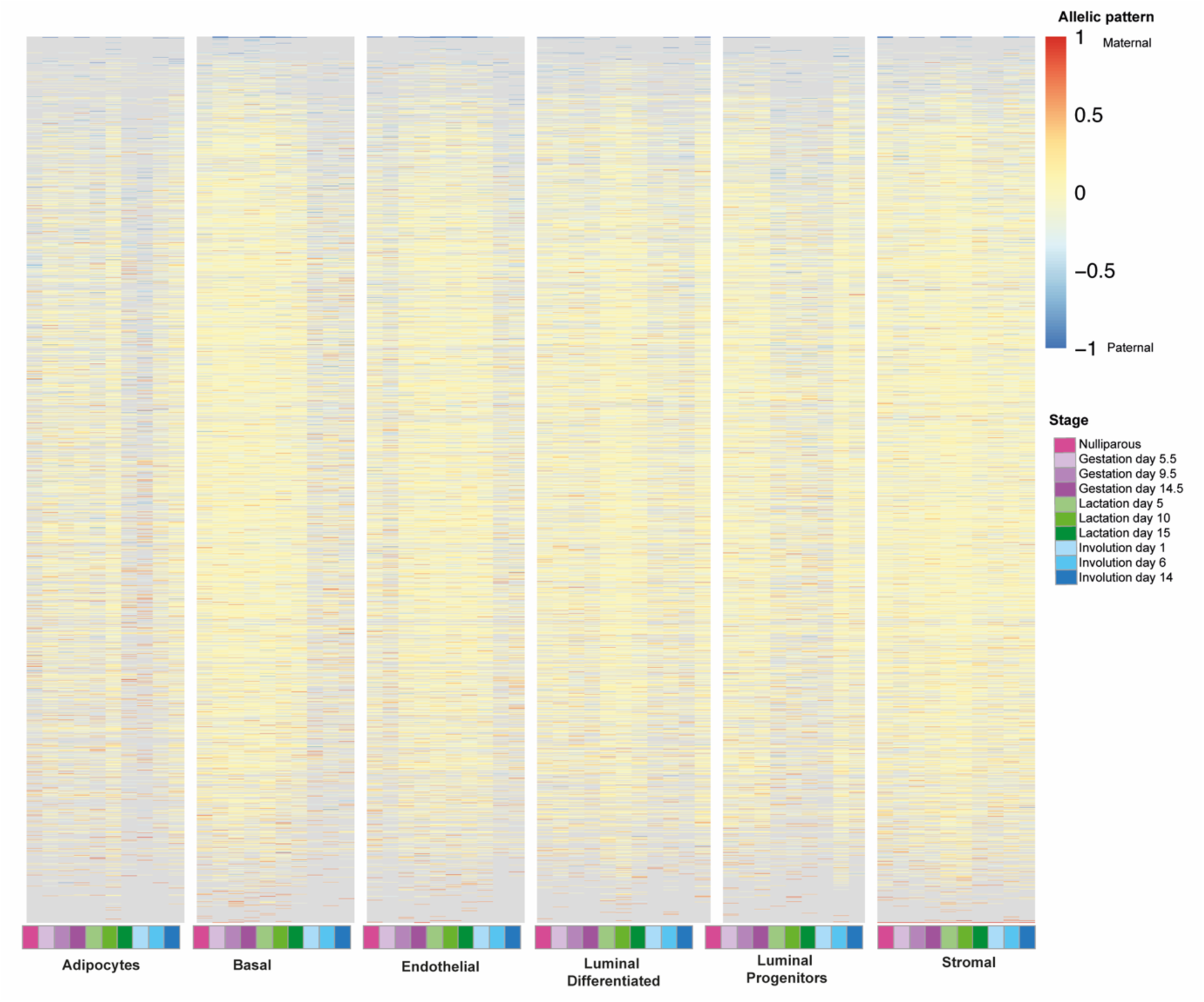
Heat map of allelic pattern of all genes expressed above a threshold of RPKM>1 in at least one cell type across the adult mammary gland developmental cycle, excluding all known imprinted genes Maternal and paternal expression are indicated in red and blue respectively, while yellow indicates biallelic expression. Grey indicated an expression level lower than the threshold. Each box represents an average of 8 biological replicates, 4 of each reciprocal cross.

**Supplementary Figure 8.**
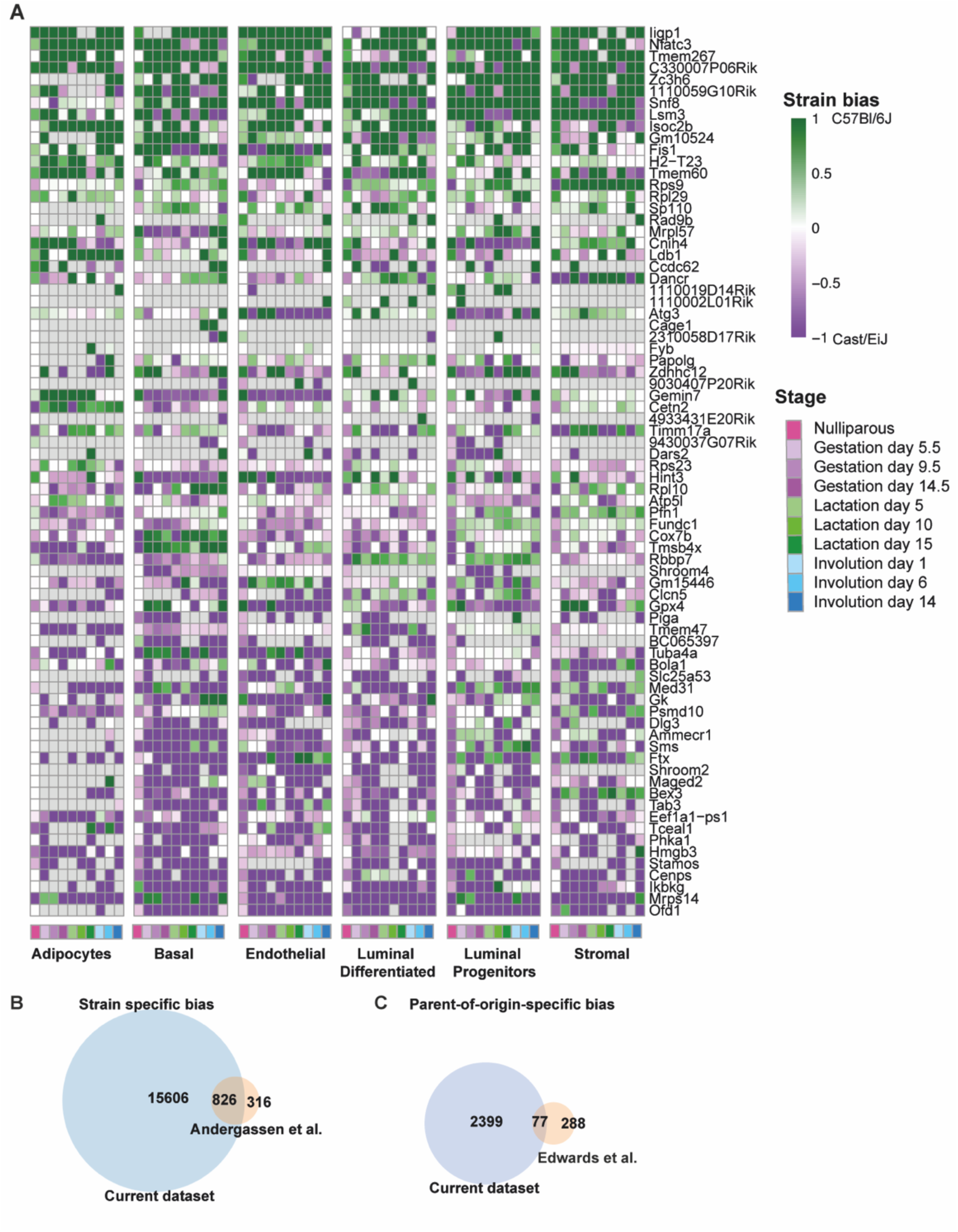
**(A)** Heat map showing statistically significant strain-specific bias between C57Bl6/J and CAST/EiJ with RPKM>1 in at least 3 timepoint or cell type as well as a bias greater than 0.5. **(B-C)** Venn diagram comparing the number of genes with strain-specific bias identified in this study compared to Andergassen et al. **(B)** or parent of origin-specific bias Edwards et al. 2023 **(C)**.

