## Supplementary table 13 for "Dynamic Allelic Expression in Mouse Mammary Gland Across the Adult Developmental Cycle"

**Supplementary table 13 – Primers used in this study.**
qPCR primers

| Gene | Forward primer | Reverse Primer |
|---|---|---|
| PDGFRa | TCCTTCTACCACCTCAGCGAG | CCGGATGGTCACTCTTTAGGAAG |
| Krt5 | TCTGCCATCACCCCATCTGT | CCTCCGCCAGAACTGTAGGA |
| Krt8 | ACTCACTAGCCCTGGCTTCA | TCTTCACAACCACAGCCTTG |
| CD31 | ACGCTGGTGCTCTATGCAAG | TCAGTTGCTGCCCATTCATCA |
| Adipoq | TGTTCCTCTTAATCCTGCCCA | CCAACCTGCACAAGTTCCCT |
| β-Tubulin | TTCAGCTGACCCACTCACTG | AGACAGGGTGGCATTGTAGG |

Pyro

| Gene | Forward primer | Reverse Primer | Sequencing Primer |
|---|---|---|---|
| Cdkn1c | TAGCAGGAACCGGAGATGG | [Btn] ACACCTTGGGACCAGCGTACT | TGGAAATCTGAAAACTGT |
| Meg3 | CTCCTGGATTAGGCCAAAGC | [Btn] GGCCAGGGTCCAGAGTCTT | GACCCTCCAACTGTAAA |
| H19 | GGGGGGTAGGATATATGTATTTTT | [Btn]ACCTCATAAAACCCATAACTATAAAATCAT | GTGTGTAAAGATTAGGG |
| Dlk1 | [Btn]CGCAAGAAGAAGAACCTCCTGT | ACGCTGCTTAGATCTCCTCATCA | CAGCCTCCTTGTTGAA |
| Igf2 | TCACGTCCCACACTAAGATCTCTC | [Btn]GGGGTGTCAATTGGGTTGT | AAGGGGATCTCAGCA |
| Snrpn | TAAATCTCAGCCCTTCTCTTCCC | [Btn]AATGCAGTAAGAGGGGTCAAAAA | CCCTTCTCTTCCCCTA |
